# Ultrahigh-throughput screening for sialic acid-active enzymes in the microbial genetic diversity

**DOI:** 10.64898/2026.08.27.747595

**Authors:** John Martínez-Salvador, Sara Trujillo-Cubillo, Laura Blas-Muñoz, Melissa Conte, Wolf-Dieter Fessner, Simon Charnock, James Finnigan, Aurelio Hidalgo

## Abstract

Sialic acids (Sias) and related nonulosonic acids are critical components of glycoconjugates involved in host-pathogen interactions, immune regulation, and cell signalling. Despite their biotechnological relevance, the diversity of enzymes involved in Sia biosynthesis remains largely underexplored due to limitations in culture-dependent methods and the lack of (ultra)high-throughput screening strategies. Here, we report the development of a highly sensitive droplet-based microfluidic screening platform enabling the functional discovery of sialic acid aldolases in environmental metagenomes. The method integrates a fluorescence-coupled enzymatic cascade compatible with fluorescence-activated droplet sorting (FADS), allowing the screening of >10⁶ droplets per experiment, as well as a downstream validation strategy for the selected hits. Although some limitations were identified, the system demonstrated high sensitivity and was utilised for the screening of a metagenomic library from garden soil. During this campaign, a potential new sialic acid aldolase enzyme was identified. This work establishes a generalizable framework for measuring complex, multi-step enzymatic functions at ultrahigh throughput using coupled cascades in droplets, with direct applicability to directed evolution and activity-guided functional metagenomics.

**Highlights:**

- Enzymatic cascade assay reports sialic acid aldolase activity in droplets.
- Dual fluorescence signal enables droplet quality control and barcoding.
- Optimized lysis and FADS enable plasmid recovery after droplet sorting.
- Droplet assay enriches NanA-positive clones by over 700-fold.
- Soil metagenome screening retrieves aldolase-related candidates

**Graphical abstract:** Fig. 1.
Optimization of a fluorescent cascade assay for NanA detection in droplets, validation using a mock library, and application to functional metagenomic screening. N-acetylneurminic acid (Neu5Ac), N-acetylmannosamine (ManNAc), sialic acid aldolase (NanA), Lacatate dehydrogenase (LDH), Ketorreductase Fb (FbKRED), Fluorescence Activated Droplet Sorting (FADS). Figure created with BioRender.

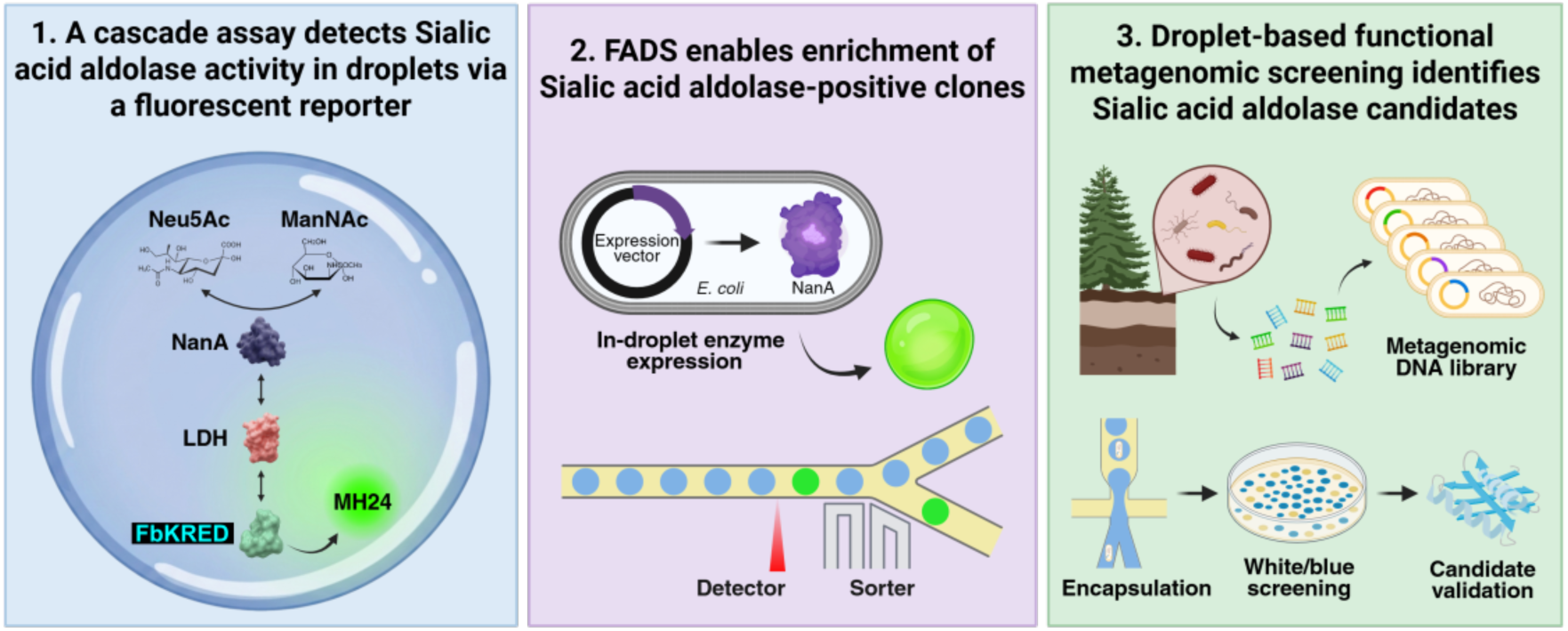

## Introduction

Nonulosonic acids, commonly referred to as sialic acids (Sias), constitute a family of nine-carbon α-keto sugars that typically occupy terminal positions in membrane glycoconjugates, with N-acetylneuraminic acid (Neu5Ac) as the most prominent member. Present across all domains of life, Sias exhibit extensive chemical and structural variability, with dozens of naturally occurring derivatives [1,2]. Due to their diversity and cellular localization, sialoconjugates play key biological roles, especially in cellular recognition and communication through specific interactions with ligands and receptors [3]. These functions are central to immune and developmental processes including regulation of the alternative complement pathway via selective recognition of sialylated surfaces [4], or control of neuronal migration and axonal growth through the neural cell adhesion molecule, a well-studied sialoconjugate highly expressed in the neural tissue [5]. Given their involvement in key biological processes, the industrial production of Sias or Sia derivatives for biotechnological applications has attracted growing interest. Applications cover functional food ingredients such as synthetic sialylated human milk oligosaccharides for infant nutrition [6], diagnostic tools, or glycoengineered therapeutics, including anticancer agents [7] and Sia analogues that inhibit viral infections by blocking interactions with host cell receptors [8].

The demand for efficient Sia production and diversification has driven interest in nature-inspired sustainable biocatalytic routes that avoid traditional multi-step chemical synthesis while enhancing stereoselectivity and atom economy [9]. For instance, bacterial sialylation processes rely on a combination of de novo biosynthesis and scavenging of host-derived sugars, followed by their incorporation into glycoconjugates. This dual strategy is enabled by a network of enzymes, including Sia synthase (e.g. NeuB from *Neisseria meningitidis*) or Sia aldolase (e.g. NanA from *Escherichia coli*). Specifically, Neu5Ac aldolase or NanA (EC 4.1.3.3), is a member of the dihydrodipicolinate synthase (DHDPS/NAL) superfamily that typically forms a homotetramer of (β/α)8-barrel subunits of approximately 33 kDa, depending on the species. It catalyses the reversible aldol cleavage of Neu5Ac releasing N-acetylmanosamine (ManNAc) and pyruvate via a class I Schiff-base mechanism, with a conserved Lys catalytic residue in a TIM-barrel scaffold [10]. Moreover, since they operate without the need for nucleotide-activated intermediates, they are suitable candidates to establish simpler and more efficient biocatalytic processes for the synthesis of Sia derivatives [1].

Metagenomics represents a cultivation-independent strategy for the discovery of novel enzymes, such as NanA homologues, in the microbial diversity. Whereas shotgun metagenomics infers a targeted enzyme function from sequence, functional metagenomics identifies target metagenomic sequences via the detection of the specific biochemical activity they encode. Consequently, the key bottleneck in the discovery of suitable biocatalysts by functional metagenomics remains assay scalability and specificity [11,12]. Traditional microtiter plate workflows, which typically screen 104-105 clones per day, have become slow, resource- and labour-intensive considering the required library sizes. In contrast, ultrahigh-throughput screening (uHTS) allows the interrogation of 106 clones per day while maintaining the genotype-phenotype linkage at the single-clone level [13]. Droplet microfluidics has emerged as a uHTS-enabling technology that enables the compartmentalisation of single cells or cell-free reactions in picoliter droplets, and fluorescence-activated droplet sorting (FADS) at kilohertz rates, drastically expanding accessibility to metagenomes [14].

However, many carbohydrate transformations, including Sia synthesis and cleavage, are difficult to screen at uHT since native substrates or products are optically undetectable in the visible region. Alternatively, fluorogenic analogues can alter substrate recognition or introduce bias in the discovered enzymes [15]. A general solution is to assemble coupled enzymatic cascades that translate a primary transformation into a fluorescent signal downstream, preserving native substrates while amplifying the signal. This screening strategy succeeded, for instance, in the discovery of industrially relevant enzymes such as glycoside hydrolases, β-glucuronidases, or KREDs [14–16]. Therefore, in this work, we developed a modular cascade assay that converts Sia aldolase activity into a droplet-sortable fluorescent signal. Then, we optimized the cascade in solution and engineered a robust droplet-workflow addressing droplet generation, linearity and sensitivity, fluorophore leakage, single-cell encapsulation conditions, in-droplet growth, induction of gene expression and cell lysis. This method was later used to screen a soil metagenomic library to identify Sia aldolase-related activities.

## Materials and methods

### Materials

All materials used in this study are described in **Supplementary Table S 1** (plasmids), **Supplementary Table S 2** (oligonucleotides), **Supplementary Table S 3** (*E. coli* strains), **Supplementary Table S 4** (culture media and solutions), **Supplementary Table S 5** (key reagents). Routine bacterial cultivation conditions are described in Supplementary Methods.

### Screening cascade development

The Sia aldolase screening cascade was first assembled with purified enzymes by sequentially coupling the cleavage of Neu5-Ac by *E. coli* NanA to the reduction of the released pyruvate by L-lactate dehydrogenase (LDH, EC 1.1.1.27) [19] and finally, to the ketoreductase (KRED)-catalysed recycling of the nicotinamide cofactor and the oxidation of the non-fluorescent alcohol MH32 (4-[6-(1-hydroxyethyl)naphthalen-2-yl]-1,1-dimethylpiperazin-1-ium iodide) to the fluorescent ketone MH24 (4-[6-acetylnaphthalen-2-yl]-1,1-dimethylpiperazin-1-ium iodide) as described by Blas-Muñoz *et al.* [18].

LDH from rabbit muscle was purchased from Merck and used as received. NanA was recombinantly produced in *E. coli* BL21 cells harbouring a pkk-233 plasmid encoding the *NanA* gene with a terminal His-tag. Cells were grown at 37 °C in Lysogeny Broth (LB) until reaching an OD_600_ of 0.4-0.6. Protein expression was induced by addition of IPTG (0.5 mM), and cultures were incubated overnight at 30 °C. Cells were harvested by centrifugation, washed, and disrupted using a high-pressure homogenizer (Panda2000, GEA, Italy). After removal of cell debris by centrifugation, the supernatant was collected and lyophilized for storage. The fluorogenic module employed FbKRED, a ketoreductase identified by screening a commercial collection of available KREDs (Prozomix Ltd., UK) using the oxidation of the fluorogenic alcohol mentioned above. When required, NanA or FbKRED were purified from lyophilized extracts by immobilized metal affinity chromatography using a Ni^2+^-charged gravity-flow column and eluted with 250 mM imidazole.

Increasing amounts of enzyme activity units were used along the cascade to ensure that the auxiliary reactions did not become rate-limiting relative to the main NanA reaction. Specific activities were determined from initial reaction rates measured using a CLARIOstar® microplate reader (BMG Labtech, Germnay) at 30 °C and reported in units (U) per milligram of protein (mg^-1^). One U is defined as the amount of enzyme catalysing the conversion of 1 µmol of substrate per minute.

For optimisation purposes, LDH and NanA activities were assayed as a single and coupled reactions, respectively, by following NADH consumption at 340 nm in 96-well microplates. Activities were calculated from the initial reaction rates using the Lambert-Beer law and an experimentally determined extinction coefficient (ε = 1.70 mM^-1^). FbKRED activity was determined fluorometrically in black, flat 384-well microplates using excitation and emission wavelengths of 340 ± 10 and 520 ± 10 nm, respectively. Relative fluorescent units (RFUs) were converted into concentration values using an experimentally determined calibration factor (1.75·10^7^ RFU mM^-1^) The final composition of the optimized screening cascade is summarized in **Table 1**.

**Table 1.** Reaction mixture of screening assay. N-acetylneuraminic acid (Neu5Ac), Lactate Dehydrogenase (LDH).

| Component | Final concentration (or unit) |
| --- | --- |
| Neu5Ac | 2.5 mM |
| NADH | 2.5 mM |
| MH32 | 1 mM |
| LDH | 0.02 U |
| FbKRED | 0.025 U |
| TRIS buffer pH 7.5 | 50 mM |

### Microfluidic device fabrication and operation

Microfluidic devices for droplet generation, picoinjection and FADS were designed in Draftsight CAD (Dassault Systèmes) and fabricated in-house by standard soft lithography using PDMS replicas cast from SU-8 master moulds [14]. Detailed device layouts are shown in **Supplementary Fig. S 1** and fabrication procedures are provided in the Supplementary Methods and **Supplementary Fig. S 2.**

Three complementary droplet generation and manipulation approaches were employed throughout this study. Monodisperse 20 and 30 μm w/o emulsions were generated using either flow-focusing or co-flow microfluidic devices. Co-flow devices were exclusively used to generate the binary control droplet mixture, whereas in all other experimental applications, flow-focusing devices were used. Droplet formation was achieved using HFE-7500 fluorinated oil containing 1.5% (w/w) RAN 008-FluoroSurfactant (RAN Biotechnologies) as the continuous phase, while precision syringe pumps (NemeSYS S, Cetoni GmbH, Germnay) were used to control flow rates. Droplet size was adjusted by modifying the relative flow rates of the continuous and dispersed phases.

Picoinjection was performed using a dedicated microfluidic device to introduce the screening assay reagents and cell lysis components into pre-formed monodisperse droplets during the metagenomic screening campaign. Injection was achieved by electrocoalescence under an externally applied electric field, allowing uniform reagent delivery into individual droplets [20].

All droplet populations were analysed or sorted using a custom FADS platform based on a design from Mazutis *et al* [14]. During sorting experiments, fluorescence signals from individual droplets were used to trigger dielectrophoretic sorting, enabling the isolation of putative NanA-positive droplets. The FADS platform consisted of a fibre-coupled multi-wavelength laser module (CNI Laser, Changchung, China) and an inverted fluorescence microscope (DMi8, Leica Microsystems, Germany) equipped with a Phantom VEO 1310L high-speed camera (Vision Research, USA). Picoinjection was performed by electrocoalescence to introduce the reaction mixture into pre-formed droplets. During sorting experiments, fluorescence signals from individual droplets were used to trigger dielectrophoretic deflection, enabling the isolation of putative NanA-positive droplets. Detailed operating conditions and parameters of droplet generation, picoinjection, analysis and sorting are provided in the Supplementary Methods.

### Establishment of the droplet-based screening assay

The assay was adapted to a droplet-based format by evaluating the stability, sensitivity and in-droplet cell lysis under screening conditions. We evaluated in-droplet stability of the fluorophore through a leakage assay. Two water-in-oil (w/o) emulsions containing the complete screening reaction mixture with LB medium and IPTG were generated, one emulsion supplemented with 0.75 mM fluorescent ketone MH24 and the other lacking the fluorophore. The emulsions were mixed at a 1:10 ratio and incubated statically at 30 °C for 72 h. At 0, 24, 48 and 72 h, droplet fluorescence was qualitatively assessed by fluorescence microscopy using an Olympus BX50 microscope equipped with an FITC filter set, a ×25 objective and a Pike F032B camera (50 ms exposure; Allied Vision, Germany). Transport of ketone MH24 between droplets was quantified by analysing emulsion samples using a fibre-coupled multi-wavelength laser module (CNI Laser, Changchung, China) with 405 nm excitation.

The sensitivity of the system was evaluated by generating w/o emulsions containing increasing concentrations (0, 0.05, 0.1, 0.5, 0.75 and 1 mM) of MH24. Emulsions were mixed in equal proportions and analysed using the laser module described above. Texas Red™-conjugated dextran (70 kDa, TexRed) was incorporated as a fluorescent reference dye to identify droplets and distinguish those with the expected volume during FADS. Similarly, emulsions containing increasing TexRed concentrations (0, 0.05, 0.1, 0.25, 0.5, 0.75 mg mL^-1^) were analysed under identical conditions. For both fluorophores, the median fluorescence intensity of each droplet population was used to construct a calibration curve.

The limit of detection (LOD) and limit of quantification (LOQ) were determined according to Armbruster, D. A *et al*. [21] from the linear regression of the calibration curve of both fluorophores using equations (1) and (2), where *a* is the intercept, *b* is the slope, *S_a_^2^* is the variance of the regression and *S_b_^2^* is the variance of the slope.

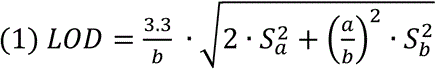

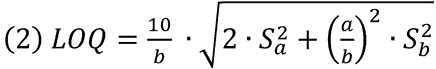

In-droplet cell lysis was evaluated by flow cytometry. Overnight cultures of *E. coli* DH10B were harvested, resuspended in 50 mM Tris-HCl (pH 7.5) and adjusted to 10⁷ cells mL^-1^. Four lysis treatments were tested: 2% (v/v) DMSO, 8 mg mL^-1^ polymyxin B, 4 mg mL^-1^ lysozyme, and a combination of polymyxin B and lysozyme. Untreated cells and cells subjected to three freeze– thaw cycles were included as negative and positive controls for cell lysis, respectively. Following incubation at 37 °C for 1 h, cells were stained with Hoechst 33342 (H33342, 1 μg mL^-1^) and propidium iodide (PI, 10 μg mL^-1^) for 15 min at room temperature in the dark and analysed using a BD FACSCanto™ II flow cytometer (BD Biosciences, USA). H33342 and PI fluorescence were detected in the FL7 and FL3 channels, respectively.

The complete screening cascade was validated in droplets using purified enzymes. Two 30 μm w/o droplet populations, corresponding to the expected droplet size after picoinjection, were generated containing the reaction mixture. Purified NanA was included to the positive population, whereas NanA was replaced with Milli-Q water in the negative population. Both populations were mixed at a 1:1 ratio, incubated under screening conditions and analysed after 0, 24 and 48 h using the microfluidic fluorescence detection system described before.

### Soil metagenomic library screening and hit validation

A previously constructed and described metagenomic library from garden soil (ABSCH-IRCC-ES-258964-1, authorization number ESNC107) by Blas-Muñoz *et al.* [18] was used for functional screening. The library comprised approximately 1.5·10^6^ *E. coli* DH10B clones carrying environmental DNA fragments with an average insert size of 1.1 kbp cloned into the pBluescript SK(+) vector.

For screening campaign, the metagenomic library was diluted to achieve non-deterministic, single-cell encapsulation, described by the Poisson distribution, and encapsulated together with IPTG in 20 μm w/o droplets to induce recombinant protein expression. Following overnight incubation, droplets were picoinjected with the screening reaction mixture, lysing agents and TexRed, and incubated overnight at 30 °C to allow enzymatic turnover and signal development. Finally, droplets were sorted by FADS, and positive droplets were recovered for downstream DNA recovery and hit validation.

Following FADS, the sorted droplets were de-emulsified using 1H,1H,2H,2H-perfluoro-1-octanol (PFO, Merck). An equal volume of extraction buffer supplemented with proteinase K (1 mg mL^-1^, Sigma-Aldrich) was added, and samples were incubated for 1 h at room temperature to promote emulsion breaking, protein digestion and DNA release. Following phase separation by centrifugation, the aqueous phase was recovered, and DNA was purified using the DNA Clean & Concentrator™-5 kit (Zymo Research). Purified DNA was directly transformed into electrocompetent *E. coli* DH10B by electroporation.

Transformants were plated onto LB agar supplemented with ampicillin (100 μg mL^-1^), IPTG (0.5 mM) and X-Gal (40 μg mL^-1^). Recombinant clones carrying metagenomic inserts in the pBluescript SK(+) vector were identified by blue-white screening, and white colonies were selected for downstream characterization. Colony PCR was performed using vector-specific primers flanking the multiple cloning site to confirm insert presence and estimate insert size. PCR products were analysed by agarose gel electrophoresis.

Putative NanA-positive clones identified by microfluidic screening were validated using the thiobarbituric acid (TBA) assay as an orthogonal activity assay as described by Warren *et al.* [22]. Clarified cell-free extracts from induced metagenomic clones were incubated with the reaction mixture (1 mM Neu5Ac in Tris 50 mM pH 7.5), and residual Neu5Ac was quantified by periodate oxidation followed by TBA colorimetric detection **Supplementary Fig. S 3**. Absorbance was measured at 549 nm to determine the amount of residual Neu5Ac. Since Sia aldolase activity results in Neu5Ac consumption and consequently lower absorbance values, TBA assay results are presented as relative activity to facilitate data interpretation.

### Bioinformatic tools

Metagenomic ORFs were predicted using SnapGene (v8.0.3). Protein sequence annotation was performed using BLASTP against NCBI databases to assign putative taxonomic and functional identities. Functional assignment of candidate enzymes was further performed using InterProScan and Phyre2 through the identification of conserved domains and protein families. Three-dimensional protein structures were predicted using AlphaFold2, and model confidence was assessed based on the predicted Local Distance Difference Test (pLDDT) scores. Predicted structures were visualized and analysed using UCSF Chimera. Additionally, potential intrinsic σ⁷⁰-dependent promoters within metagenomic inserts were identified using BPROM (SoftBerry) by examining upstream regions of candidate ORFs.

## Results and Discussion

### Assembly of the coupled screening assay

Initially, a fluorescence-based screening assay was developed, in which Sia aldolase activity was coupled to a downstream fluorescent readout via a three-enzyme cascade (**Fig. 2**). The cascade reaction comprises (i) Neu5Ac (**1**) cleavage to ManNAc (**2**) and pyruvate (**4**); (ii) pyruvate-dependent NADH oxidation to lactate (**3**) by lactate dehydrogenase (LDH) and (iii) a fluorogenic KRED module that regenerates NADH while converting the non-fluorescent alcohol MH32 (**6**) into the fluorescent ketone MH24 (**5**).

**Fig. 2.**
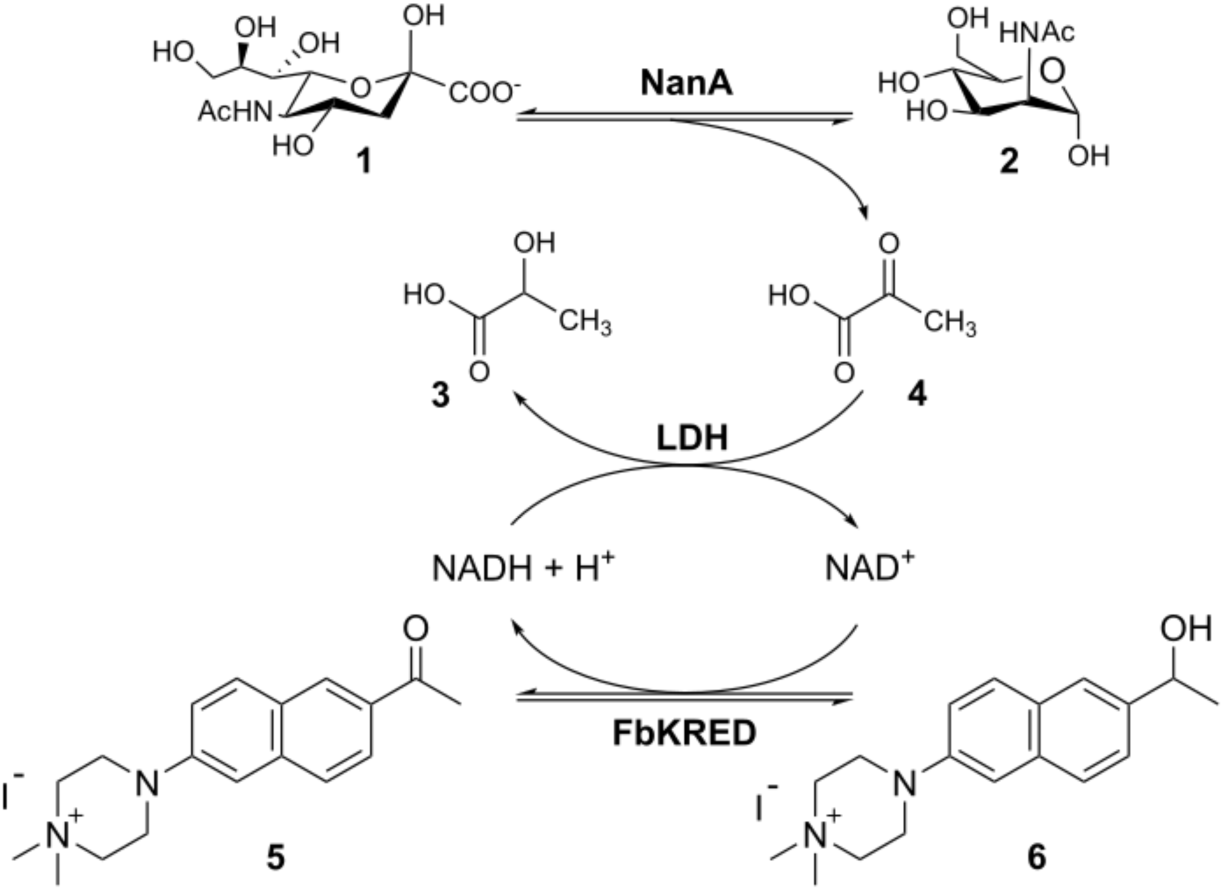
Coupled assay for Sia aldolase activity detection. The recycling assay links pyruvate generation by NanA from Neu5Ac to NADH oxidation via LDH and finally a KRED module converts the fluorogenic alcohol MH32 to the fluorescent ketone MH24. (1) N-acetylneuraminic acid, (2) N-acetyl mannosamine, (3) lactate, (4) pyruvate, (5) MH24, (6) MH32.

Since detection of the target activity by FADS required a fluorescent output, first a suitable KRED capable of specifically converting alcohol **6** into the fluorescent ketone **5** had to be identified. To that end, a commercial KRED collection was screened against alcohol **6**. The 7 best-performing candidates were subsequently coupled with the screening assay cascade to assess their signal-to-background ratio (**Supplementary Fig. S 4**). FbKRED exhibited the highest performance, by showing a high conversion of the fluorogenic alcohol **6**, while maintaining excellent substrate-specificity, thereby avoiding undesired side reactions with other cascade components or cell metabolites (**Supplementary Fig. S 4H**). Together, these properties provided the highest signal-to-background ratio, maximizing the sensitivity of the screening assay. Therefore, FbKRED was selected as the reporter enzyme.

Then, the cascade was assembled stepwise by first validating the NanA and LDH reactions through UV/Visible measurements and subsequently integrating the fluorescence-generating step, with enzyme concentrations optimised to ensure balanced catalytic flux and to avoid rate limitations. To that end, the specific activities of NanA, LDH and FbKRED were determined to be 2.01 ± 1.05, 398.84 ± 86.07 and 3.67 ± 2.20 U mg^-1^, respectively. Under selected assay conditions, the reaction containing purified NanA showed a clear increase in fluorescence compared to control reaction lacking the aldolase (**Fig. 3**¡Error! No se encuentra el origen de la referencia.**A**). These results demonstrated that the cascade efficiently coupled the Sia aldolase activity to the fluorogenic reaction, providing a robust fluorescent readout. The modular detection strategy could potentially be adapted to other complex enzymatic activities generating NADH. Additionally, the cascade was also evaluated using cell-lysates expressing NanA at concentrations equivalent to those of a single *E. coli* clone in a 20 μm droplet. The fluorescent signal using lysates remained sensitive and specific, (**Fig. 3**¡Error! No se encuentra el origen de la referencia.**B**) confirming successful enzyme coupling and assay functionality in a biologically relevant setup.

**Fig. 3.**
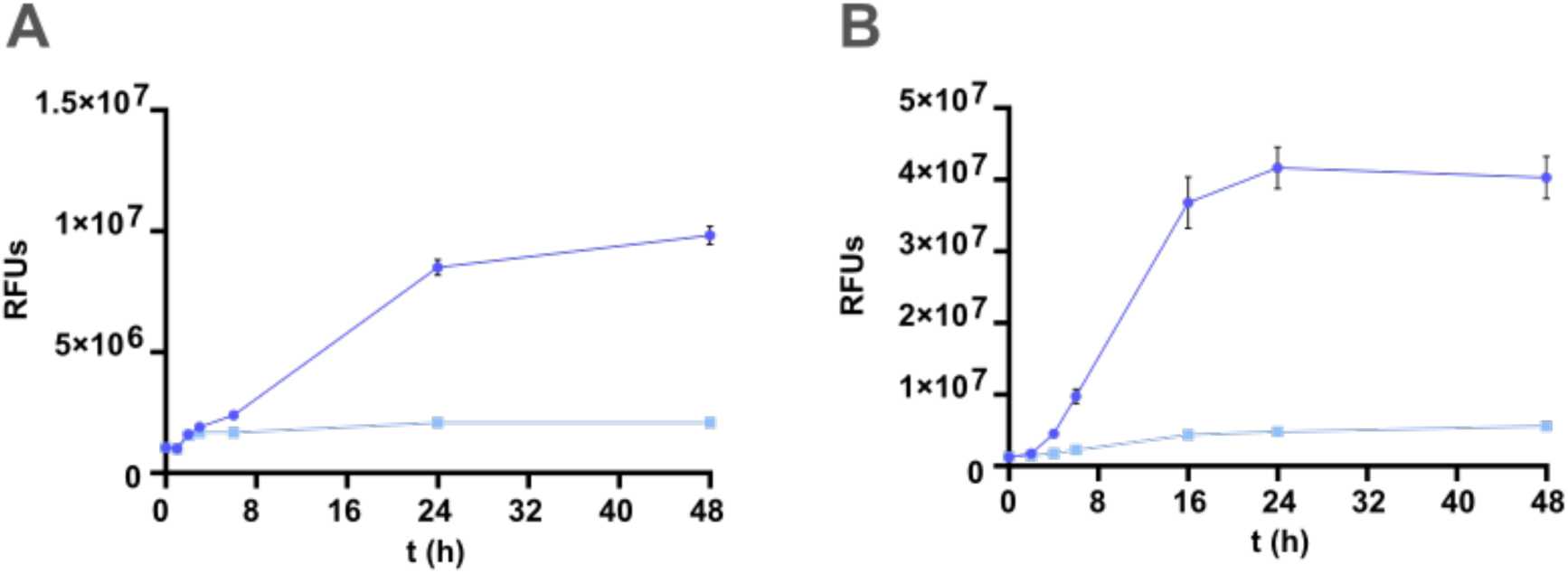
Time courses of Sia aldolase activity via a coupled assay using pure NanA (A) or an amount of cell lysate equivalent to a single cell lysed in a 20 μm droplets (B). Dark traces in black represent NanA-containing reactions while light traces in grey show negative controls lacking NanA. Data represent mean ± SD (n = 3).

### Setup of the screening assay in microfluidic droplets

To establish the assay in droplets, several key biological and analytical parameters were optimised. First, fluorophore leakage was examined using two populations of 30 μm w/o droplets containing either 0.75 mM ketone **5** or 50 mM Tris buffer. Both populations were mixed at a 1:10 ratio to mimic a screening-relevant scenario with a lower abundance of positive droplets than negative droplets. As shown in **Fig. 4**, both populations remained clearly distinguishable up to 72 h incubation, indicating very low fluorophore leakage under selected conditions and in agreement with the design of the alcohol **6**/ ketone **5** reporter pair, in which the formal positive charge increases aqueous-phase retention, as previously described [18,23]. This retention window is also consistent with reported incubation times for metagenomic screening campaigns [20] and it is sufficient for development of Sia aldolase activity signal. Thus, fluorophore leakage was not expected to compromise population discrimination during the actual sorting workflow. Nevertheless, reporter retention remains an assay-defining parameter in droplet microfluidics, and should be re-evaluated if incubation times, oil/surfactant formulations or reporter structures are modified [21–22].

**Fig. 4.**
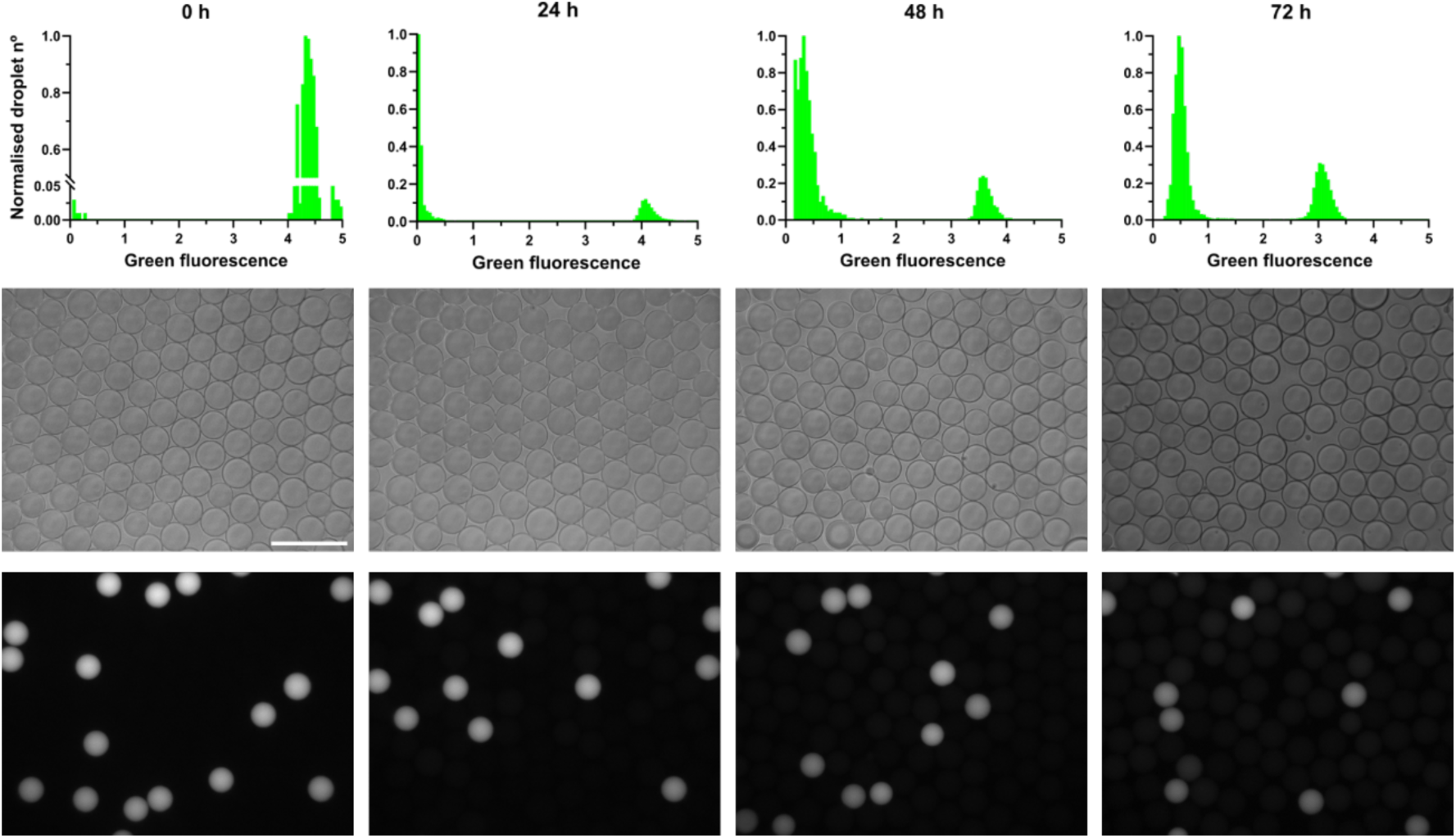
Leaking of fluorescent ketone 5 from droplets. Upper row: Green fluorescence (V) histograms at 4 time points. Middle row: Phase contrast images of a 1:10 mixture of droplets with and without ketone 5 after 0, 24, 48 and 72 h. Lower row: Fluorescence microscopy images at 50 ms of exposure of a 1:10 mixture of droplets with and without ketone 5 after 0, 24, 48 and 72 h. Images were analysed using Fiji image processing software (I). Scale 100 μm.

Then, the sensitivity of the two fluorophores needed for the droplet screening (ketone **5** as the product reporter and TexRed, as droplet volume and identity tracer) was determined (**Fig. 5**). Calibration curves were generated by plotting the median droplet fluorescent intensity against fluorophore concentration under assay conditions. The highest green fluorescence signal (1 mM), equivalent to a complete conversion of alcohol **6** into ketone **5**, was detected at 6 V, while TexRed concentrations used in this study (0.1 and 0.25 mg mL^-1^) were detected at 1.10 V and 2.65 V respectively. This result demonstrated that the activity reporter concentrations used in the screening assay, as well as the tracer concentrations selected, are correctly detected within the linear range of the FADS platform. The TexRed signal was used as an internal droplet-quality criterion to improve event classification and sorting reliability by excluding droplets displaying abnormal fluorescence resulting from coalescence, incomplete picoinjection or volume heterogeneity. This dual-fluorophore strategy enabled activity-dependent fluorescence to be assessed independently of droplet volume variations and allowed reliable gating of the main droplet population. This distinction is particularly relevant for FADS, where fluorescence intensity is used to classify and sort individual droplets and technical variability could otherwise be misclassified as differences in enzymatic activity.

**Fig. 5.**
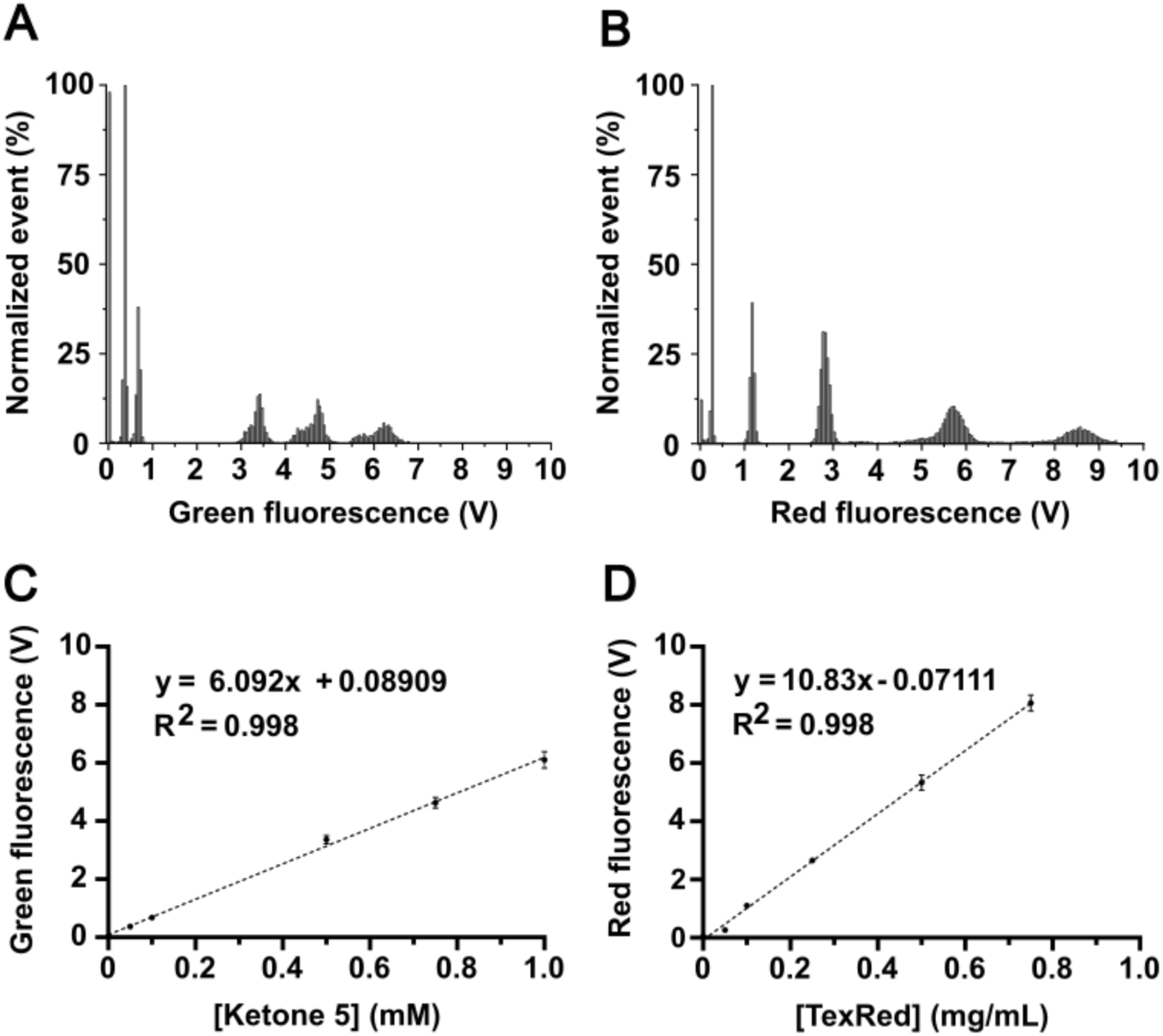
Calibration curves of fluorophores MH24 and TexRed in microfluidic droplets. Histograms showing normalized numbers of droplets containing increasing concentrations of MH24 (A) and TexRed (B), whose median value was represented against the concentration of MH24 (C) and TexRed (D).

The analytical sensitivity of both fluorescence channels further supported this dual-fluorophore strategy. Ketone **5** showed a LOD of 58 μM and LOQ of 175 μM while TexRed showed a LOD of 31 μM and LOQ of 96 μM. These detection and quantification limits indicate that small changes in reporter concentration can be reliably detected under the conditions used for droplet screening. Overall, these parameters established the detection conditions used during a metagenomic screening campaign.

To maximise the release of intracellularly produced enzymes in droplets, different methods for cell lysis were tested via cell membrane integrity using dual H33342/PI cell staining and flow cytometry, where H33342 labels all cells and PI only stains membrane-compromised cells [25–26]. The determination of cell integrity was calibrated using defined mixtures of viable and freeze-thaw-treated *E. coli* cells, showing a concomitant increase in membrane-disrupted events (PI-positive) (**Supplementary Fig. S 5**). Then, four chemical lysis treatments were evaluated under assay conditions. While 2% DMSO and lysozyme alone resulted in minimal permeabilization (<5% PI-positive cells), polymyxin B increased this fraction to 14.45%. The combined treatment with polymyxin B and lysozyme yielded the highest permeabilization (30.7% PI-positive cells), identifying it as the most effective condition for enzyme release (**Supplementary Fig. S 6**). This result aligns with previous studies where the polymyxin B and lysozyme combination enabled enzyme release prior to FADS [29]. Efficient cell lysis would maximise the amount of available NanA or metagenomic enzymes that can trigger the cascade, thereby increasing reporter accumulation and assay sensitivity, particularly for low-activity enzymes, enzymes requiring cofactors or correct folding environments [30]. Furthermore, improved lysis should facilitate plasmid release and recovery following droplet sorting, enhancing the efficiency of downstream hit identification.

Finally, the assay was evaluated in 30 μm w/o droplets by assembling the screening cascade as previously established to ensure a balanced catalytic flux. The positive droplet population contained purified NanA, whereas in the negative population NanA was replaced with Milli-Q water. Positive and negative emulsions were mixed at a 1:1 ratio and analysed at 0, 24 and 48 h by FADS. As shown in **Fig. 6**, at 0 h, a signal separation of 1.40 V was observed, increasing to a maximum of 2.75 V at 24 h. After 48 h, signal separation decreased to 2.45 V, suggesting that the assay had reached its useful dynamic range under droplet conditions. Overall, the screening assay exhibited rapid signal development, demonstrating that the cascade is functional under droplet conditions, successfully coupling Sia aldolase activity to a fluorescence readout suitable for FADS-based metagenomic screening. Additionally, the clear separation between the two droplet populations proved the robustness of the assay for fluorescence-based discrimination. Therefore, a 24 h incubation period was selected as the optimal compromise between signal intensity, fluorophore leakage and droplet stability, since extending the incubation to 48 h did not further improve positive and negative separation.

**Fig. 6.**
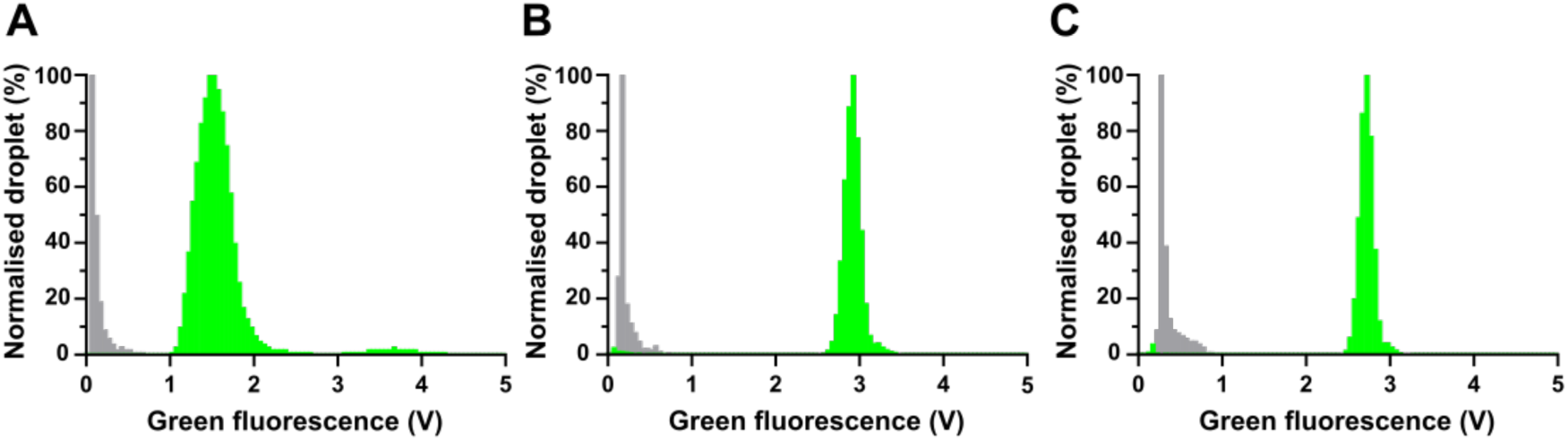
Fluorescent signal histograms showing 1:1 mixture of positive to negative droplet populations after 0 h (A), 24 (h) and 48 (h) of incubation.

A distinctive feature of this assay is the length of the enzymatic cascade that can be operated within picoliter droplets. Previous droplet-compatible coupled assays have elegantly addressed the problem of optically silent substrates by connecting the target activity to common detectable intermediates, such as carbohydrate products, H₂O₂ or NAD(P)H [17,31] [32]. However, these systems generally rely on one or two auxiliary detection steps downstream of the target reaction. In contrast, the present NanA assay requires coordinated operation of three enzymes inside the droplet, and the same logic can be extended to NeuB by adding the upstream synthase reaction, resulting in a four-enzyme cascade. This indicates that complex, multi-enzyme biochemical logic can be preserved under FADS-compatible droplet conditions, minding the specificity of the involved enzymes.

### Enrichment capacity and DNA recovery

The enrichment capacity of the droplet screening was assessed by separating a binary droplet mixture generated by encapsulating single-cells of *E. coli* expressing NanA from a pBS plasmid under the control of the *lac* promoter (positive droplets), labelled with 0.25 mg mL^-1^ TexRed, and *E. coli* harbouring an empty pBS plasmid (negative droplets), labelled with 0.10 mg ml^-1^ TexRed. Emulsions were mixed at a 1:100 ratio, incubated off-chip overnight for signal development, and a total of 150,000 events were analysed by FADS.

As shown in **Fig. 7A**, positive and negative droplets were clearly discriminated. A total of 53 droplets included within the positive sorting gate were recovered, which represented approximately 0.035% of the total analysed events. Following de-emulsification, the recovered DNA was directly transformed into competent *E. coli* DH10B cells. Transformants were subjected to blue-white screening yielding 45 white colonies and 8 blue colonies. Colony PCR confirmed the presence of the *nanA* insert in 39 of the 53 recovered transformants, corresponding to 73.6% positives (**Fig. 7B).** This enrichment increased the fraction of enzyme-positive clones from 0.1% to 73.6%, equivalent to a >700-fold enrichment capacity over the initial abundance of positive droplets.

**Fig. 7.**
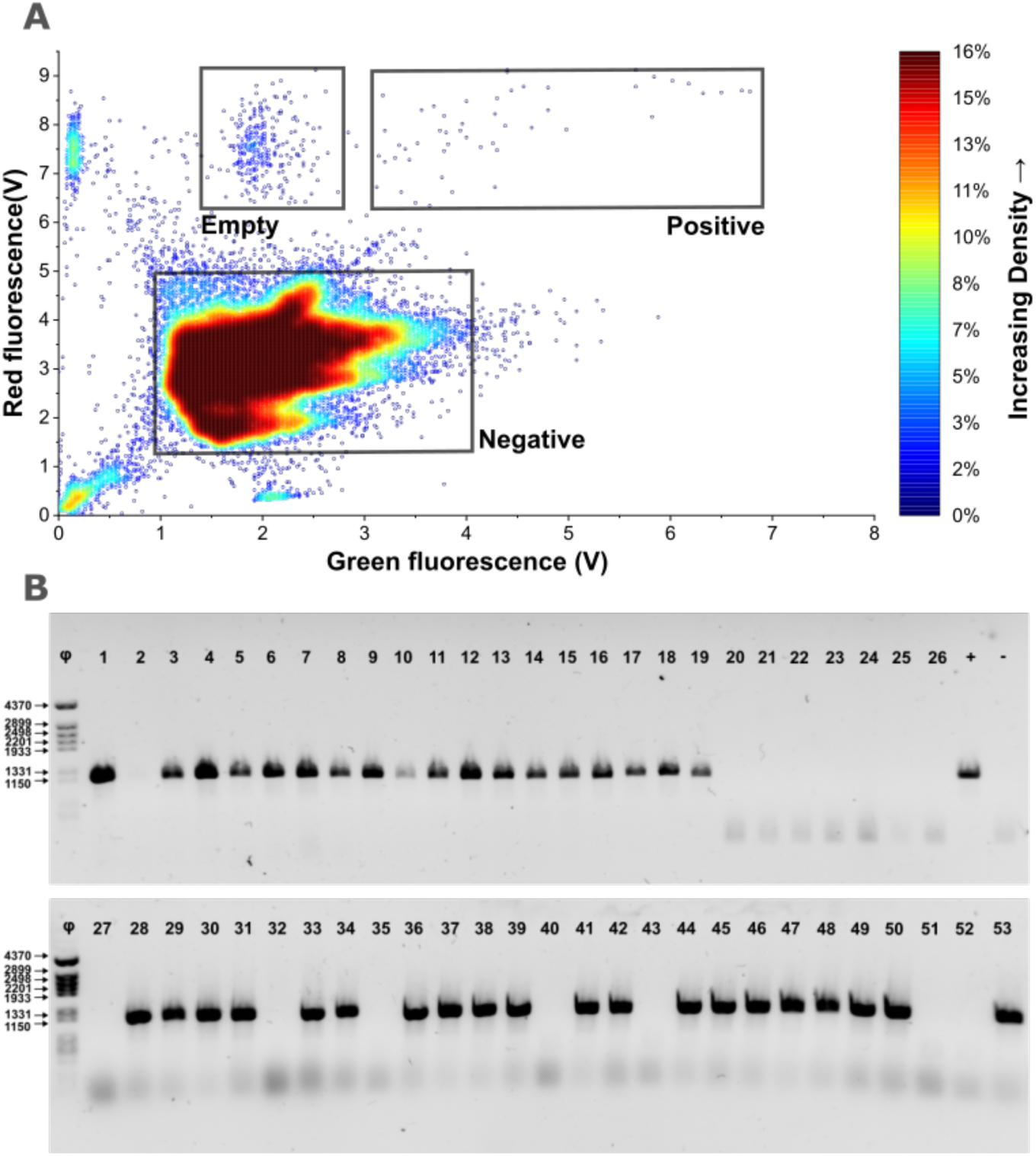
Separation of an emulsion containing a 1:100 ratio of NanA-positive to NanA-negative droplets. FADS analysis of the emulsion (A), showed populations of droplets containing NanA-expressing clones, empty droplets due to single-cell encapsulation and negative control droplets. Droplets containing NanA-expressing clones were sorted, pooled, de-emulsfied, plasmid DNA purified and analyzed by PCR using NanA-specific primers (B). PCR confirmed white colonies: 1-19, 28-31, 33, 34, 36-39, 41, 41, 44-50 and 53. PCR confirmed blue colonies: 20-26, 27, 32, 35, 40, 43, 51 and 52. “+”: nanA expressing pBS as positive control. “-”: empty pBS as negative control. Φ: DNA lader, molecular weight marker

Moreover, direct transformation of the recovered DNA enabled the reliable recovery of positive transformant clones from as few as 53 sorted droplets, demonstrating that the de-emulsification procedure preserved plasmid integrity and functionality despite the use of PFO, a de-emulsifying agent reported to interfere with certain molecular biology applications, such as PCR [33]. The direct extraction of plasmid DNA from de-emulsified droplets enabled a more straightforward recovery of genetic material over PCR-based strategies, since it avoids amplification biases and additional re-cloning steps associated with PCR-mediated recovery of sorted clones [34]. Collectively, these results validate the complete screening workflow, from clone encapsulation to genetic recovery.

### Screening of a metagenomic soil library and confirmation of hits

To illustrate the applicability of the droplet assay in a real metagenomic screening campaign, a soil-derived metagenomic library was screened to identify novel NanA homologues. This library contained 1.5 million clones and was constructed as described in Blas-Muñoz, L. *et al* [18]. A total of 4.2 million droplets were analysed, corresponding to ∼420,000 clones (assuming stochastic encapsulation of single *E. coli* cells following the Poisson distribution) and an approximate library coverage of 28%. The sorting gate for positives was set over 6 V to ensure a stringent selection of active variants, at the expense of excluding weakly active variants. This gating enabled the isolation of 2,029 droplets (0.05% of total events) (**Fig. 8**).

**Fig. 8.**
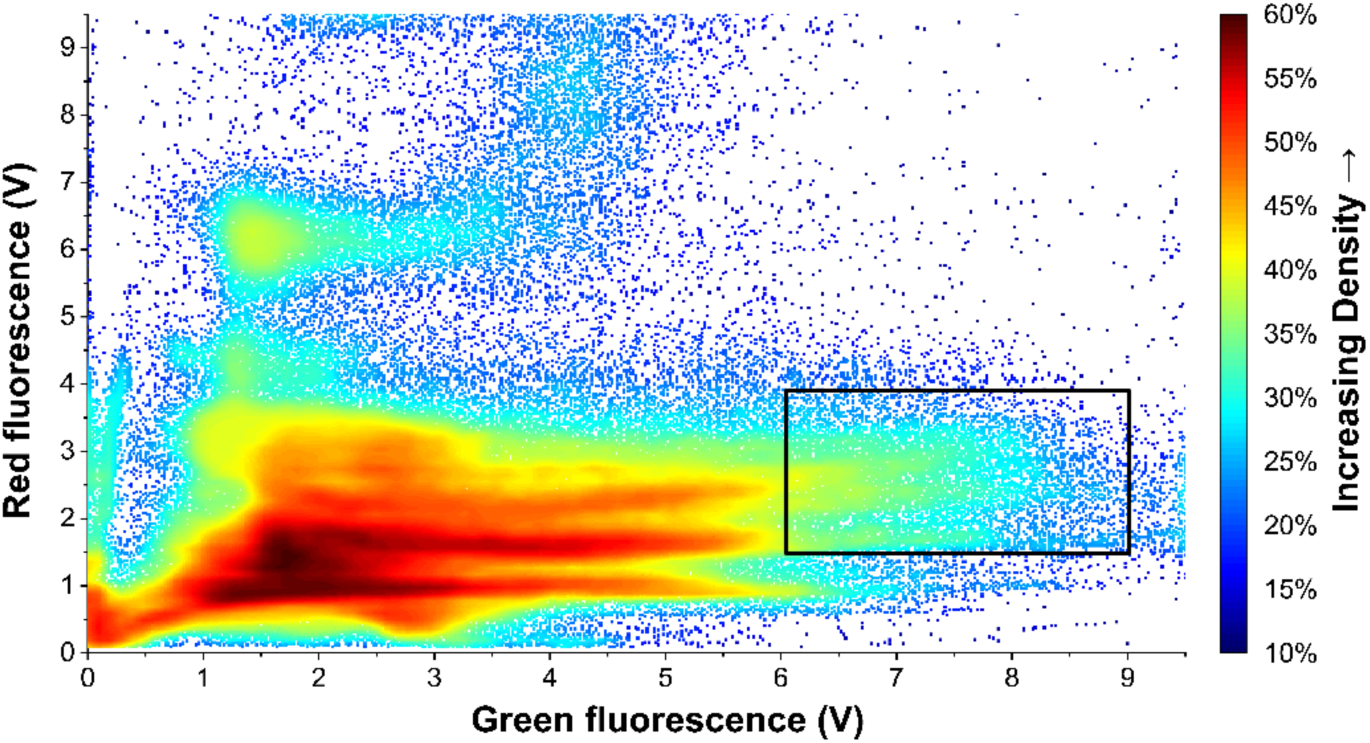
FADS dot-density plot generated for the soil metagenomic library screening campaign. Green fluorescence (x-axis) and red fluorescence (y-axis) were measured for MH24 reporter and TexRed tracker signals respectively. The gated population selected for sorting is indicated by the black rectangle (green fluorescence: 6-9 V; red fluorescence: 1.5-4 V). The colour scale represents increasing droplet density.

Recovered plasmid DNA was directly transformed into competent *E. coli* DH10B cells. Blue-white screening yielded 757 white colonies. Then, insert-containing clones were screened for activity using lysate with the aldolase-LDH coupled assay and monitoring NADH depletion stectrophotometrically. Based on a threshold defined as the negative control mean plus three times its standard deviation (5.57 μM min⁻¹), 78 clones (10.3%) were classified as active (**Supplementary Fig. S 7**).

Subsequent insert size assessment by colony PCR excluded eight clones containing metagenomic inserts shorter than 300 bp. Sequencing of the remaining candidate clones revealed that 61 of these 70 clones contained a single ORF that corresponded to the *NanA* construct used to set up the assay. The nine clones with unique inserts were further validated using the orthogonal TBA assay (**Fig. 9**), to reduce the probability of false-positive candidates arising from artefacts associated to the FADS. These candidates displayed heterogeneous activity profiles, and statistical analysis (one-way ANOVA followed by Dunnett’s test) identified two clones with significantly higher activity than the negative control (adjusted p ≤ 0.0001), whereas the remaining variants were considered non-significant.

**Fig. 9.**
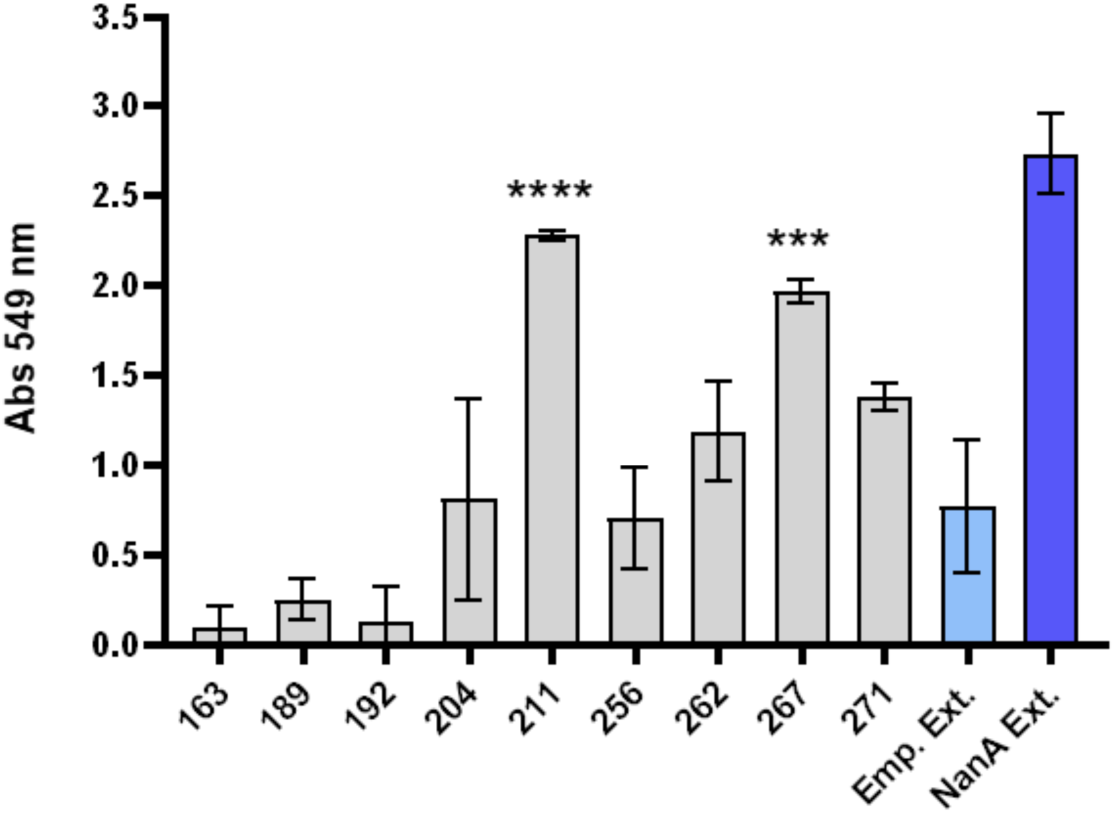
Validation of putative NanA-positive metagenomic clones using the thiobarbituric acid (TBA) assay. Bar graph showing the relative Sia aldolase activity of the nine candidate clones. Since Sia aldolase activity results in decreased absorbance at 549 nm due to Neu5Ac consumption, data are presented as relative activity so that higher bars correspond to higher enzymatic activity. Light blue indicates negative control (empty extract), while dark blue indicates positive control (NanA expressing extract). Data represent mean ± SD (n = 3). Statistical significance was determined by one-way ANOVA and Dunnett’s multiple comparison test; **** P < 0.0001, *** P = 0.0001.

The small number of recovered candidates is likely a consequence of the stringent gating applied during sorting. Such restrictive strategies preferentially select clones displaying high catalytic activity and often reduce or eliminate the need for iterative enrichment rounds. However, they may also exclude genuine metagenomic hits. Environmental genes are frequently expressed at low levels in heterologous systems due to differences in promoter recognition, codon usage or protein folding [27–28], resulting in reduced activities generating fluorescence signals below highly selective sorting thresholds. A more permissive sorting gate combined with rounds of enrichment could increase the probability of recovering low-activity clones, although at the cost of increased screening effort and a larger number of false-positive candidates requiring validation.

The limited number of validated hits may also reflect the relatively low abundance of Sia catabolic pathways in soil communities. A recent metagenomic survey demonstrated the presence of Sia genes involved in Sia catabolism in soil metagenomes, however, their frequency is typically below 10 hits per Gb and substantially lower than in host-associated microbiomes [37]. Moreover, only approximately one-third of the library was screened in the present study. Increasing library coverage would likely increase recovery rates. Nevertheless, the marked enrichment of the positive-control construct among the recovered clones demonstrates the specificity of the assay and the effectiveness of the sorting strategy for discriminating highly active variants from the metagenomic background. Additionally, the identification of two uncharacterized NanA candidates reveals the ability of the platform to recover novel functional enzymes from environmental metagenomic libraries.

### Bioinformatic analysis of putative hits

To identify the genes underlying the observed activity, metagenomic inserts from clones 211 and 267 were analysed to identify predicted open reading frames (ORFs). Clone 211 contained a 2.63 kb insert encoding eight predicted ORFs, whereas clone 267 was discarded due to the absence of candidate ORFs. Sequence similarity analysis using BLASTP assigned the insert of clone 211 to the genus *Bradyrhizobium* (100% query coverage and 90.3% sequence identity). Among the eight predicted ORFs, only three showed high similarity to proteins assigned to the genus *Bradyrhizobium* (**Supplementary Fig. S 8**). ORF 211_1 encoded a protein of 542 amino acids and showed the highest similarity to a GH17 family glycoside hydrolase (100% query coverage and 90.2% sequence identity); whereas ORF 211_5 encoded a 191 amino acid protein and matched a PLP-dependent transferase (99% query coverage and 96.8% sequence identity). ORF 211_6 encoded a 134 amino acid protein, displayed considerably lower sequence similarity to known proteins (64% query coverage and 66.3% sequence identity) and could not be assigned to any protein family based on BLASTP analysis (**Supplementary Table S 6**).

Functional annotation further supported these assignments. Consistent with the BLASTP results, InterProScan classified ORF 211_1 as a member of glycoside hydrolase family 17 (GH17), while Phyre2 predicted a GH17 β-1,3-glucanosyltransferase fold with 100% confidence, although only 48% of the sequence could be modelled. In contrast, ORF 211_5 was annotated by InterProScan as a pyridoxal 5′-phosphate (PLP)-dependent enzyme related to the L-threonine aldolase family. This annotation was strongly supported by Phyre2, which predicted an L-threonine aldolase fold with 99.8% confidence and 96% sequence coverage. ORF 211_6 lacked conserved domains and could not be functionally annotated with confidence.

Structural modelling using AlphaFold was consistent with these predictions. Both ORFs 211_1 and 211_5 produced well-defined globular models with high per-residue confidence scores (pLDDT >70 across most of the sequence), whereas lower-confidence regions were mainly restricted to terminal or loop regions. The structure predicted for ORF 211_1 (**Fig. 10**) was consistent with the characteristic fold of GH17 glycoside hydrolases, whereas ORF 211_5 displayed a partial TIM-barrel-like architecture compatible with PLP-dependent enzymes.

**Fig. 10.**
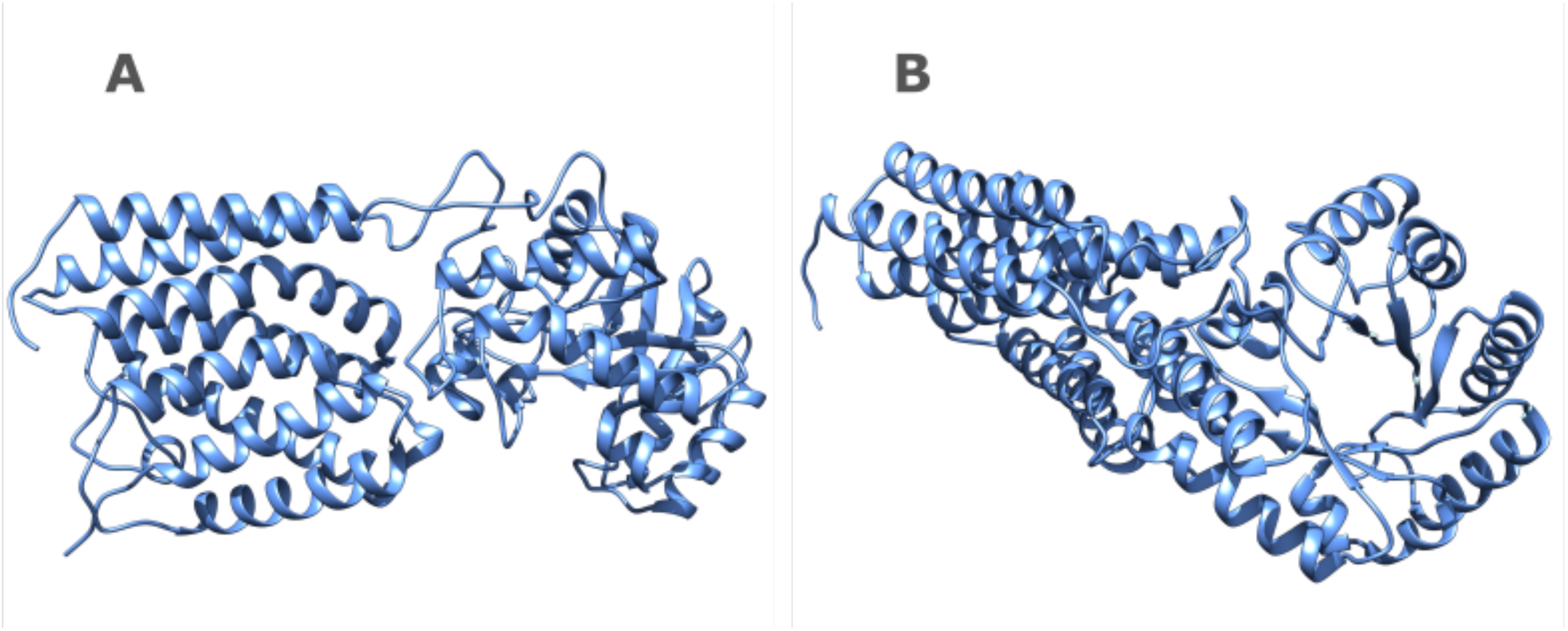
AlphaFold-predicted structure of ORF 211_1. Ribbon representation of the AlphaFold-predicted structure of ORF 211_1 shown from two different orientations. (A) Lateral view highlighting the overall two-domain architecture connected by a narrow linker. (B) Front view showing the characteristic barrel-like core of the predicted GH17 glycoside hydrolase fold. The structure is displayed as a blue cartoon representation..

Interestingly, ORFs 211_5 and 211_6 were oriented opposite to the *lac* promoter of the pBluescript vector. Expression of these genes would therefore depend on the presence of a native promoter within the metagenomic insert [38]; however, promoter prediction using BPROM failed to predict any plausible promoter sequence upstream of either ORF. In contrast, ORF 211_1 was positioned in the correct orientation for transcription from the vector promoter, making it the most plausible candidate responsible for the observed activity in clone 211.

Taken together, these analyses identify ORF 211_1 as the most likely candidate responsible for the observed activity in clone 211. Despite representing the most plausible candidate based on its orientation relative to the vector promoter, none of the identified proteins showed sequence similarity to known NanA enzymes. This may reflect the identification of a highly divergent enzyme that cannot be recognised by sequence-based annotation alone. This illustrates one of the principal advantages of activity-based functional metagenomic screening: its ability to identify candidates independently of sequence homology [11]. However, the predicted annotation of ORF 211_1 as a GH17 enzyme, highlights the need for heterologous expression and biochemical characterization of the individual ORF to validate this candidate as a Sia aldolase enzyme. Nevertheless, these findings demonstrate that the developed screening platform is capable of recovering previously uncharacterized functional enzyme candidates from environmental metagenomic libraries.

## Conclusions

We have developed and validated a droplet-compatible, coupled enzyme assay that converts sialic acid aldolase (NanA) activity into robust fluorescent readouts suitable for FADS. Assay optimization in solution established functional module balancing and enabled selection of an effective KRED reporter capable of specifically yielding fluorescent reporter MH24. Translation to droplets required systematic optimization of droplet generation, fluorophore calibration, leakage control, Poisson loading, in-droplet growth/induction, picoinjection, and cell lysis. Under optimized conditions, NanA activity exhibits fast signal development and strong droplet population separation after 24 h incubation. The metagenomic library screening simulation proved the suitability of the method for the discovery of novel enzymes, and a soil metagenomic screening campaign demonstrated its ability to identify novel NanA candidates, providing a proof of concept for activity-based enzyme discovery. Beyond NanA activity, this framework utilises a NADH/NAD^+^-dependent detection module, providing a potentially generalizable strategy for other complex enzymatic reactions. The approach may be applicable to directed evolution and activity-guided functional metagenomic screening of sialic acid-metabolizing enzymes and other multistep enzymatic activities requiring coupled detection systems.

## Acknowledgments

This work has received funding under the European Commission’s Framework Programme Horizon Europe under Grant Agreements number 101081857 (“BlueTools”) and 956631 (CC-TOP). The CBM is funded by “Centre of Excellence Severo Ochoa” Grant CEX2021-001154-S from MCIN/AEI /10.13039/501100011033 and receives institutional support by the Fundacion Ramon Areces. We thank Carlota Ruíz for assistance with the AutoCAD design and preparation of the microfluidic device schematics.

## CRediT authorship contribution statement

Martínez-Salvador, John: Investigation, Methodology, Validation, Data curation, Formal analysis, Visualization, Writing - original draft, Writing - review and editing.

Trujillo-Cubillo, Sara: Investigation, Methodology, Validation, Data curation, Formal analysis, Visualization, Writing - review and editing.

Blas-Muñoz, Laura: Investigation, Methodology, Formal analysis.

Conte, Melissa: Investigation, Methodology, Data curation, Formal analysis.

Fessner, Wolf-Dieter: Conceptualization, Funding acquisition, Review and editing.

Charnock, Simon: Formal analysis, Review and editing.

Finnigan, James: Formal analysis, Review and editing.

Hidalgo, Aurelio: Conceptualization, Funding acquisition, Project administration, Writing - review and editing, Supervision.

## Appendices

## Data availability

## Declaration of generative AI use in the manuscript preparation process

During the preparation of this work, the authors used an AI-assisted writing tool to improve language. After using this tool, the authors reviewed and edited the content as needed and take full responsibility for the content of the published article.

## Supporting information

## Supplementary Methods

### Microfluidic chip design and fabrication

Microfluidic devices were designed in AutoCAD (Autodesk, USA) and fabricated in-house by standard soft lithography as previously described by Mazutis *et al*. [14]. Four device geometries were used throughout this study: flow-focusing, co-flow, picoinjection, and fluorescence-activated droplet sorting (FADS) devices. Device layouts are shown in **Supplementary Fig. S 1**.

Master moulds were fabricated by patterning SU-8 negative photoresist (MicroChem, USA) on silicon wafers using standard photolithography. Polydimethylsiloxane (PDMS; Sylgard™ 184, Dow Corning) elastomer and curing agent were mixed at a 10:1 (w/w) ratio, degassed, poured onto the master moulds (Tekniker, Spain) and cured overnight at 65 °C. After curing, PDMS replicas were peeled from the moulds and inlet and outlet ports were created using a 1 mm biopsy punch (Kai Industries, Japan).

PDMS devices were irreversibly bonded to glass slides by oxygen plasma treatment using a FEMTO plasma etcher (Diener Electronic, Germany) operated at 150 W for 20 s. Channel surfaces were rendered hydrophobic by flushing with 1% (v/v) trichloro-(1H,1H,2H,2H-perfluorooctyl)-silane diluted in HFE-7500 fluorinated oil. Devices were stored at room temperature in a dust-free environment until use.

### Monodisperse water-in-oil (w/o) emulsion generation

Unless otherwise stated, all microfluidic devices were operated using neMESYS Base 120 syringe pump modules (CETONI GmbH, Germany) controlled through the neMESYS software. Fluids were delivered using glass syringes (SGE, Australia) connected via polyethylene (PE) tubing (2Biol, ID 0.38 mm, OD 1.09 mm). The oil phase consisted of HFE-7500 fluorinated oil containing 1.5% (v/v) FluoroSurfactant (RAN Biotechnologies, USA), unless otherwise specified.

Monodisperse w/o droplets with diameters of 20 or 30 μm were generated using either flow-focusing or co-flow microfluidic devices. Flow-focusing devices contained one aqueous inlet and one oil inlet, whereas co-flow devices incorporated two aqueous inlets to enable simultaneous encapsulation of two independent aqueous streams. Droplet formation was monitored using an inverted microscope (XDS-1R, OPTIKA, Italy) equipped with a FASTCAM Mini UX50 high-speed camera (Photron, Japan).

The dispersed phase contained the biological or chemical sample of interest. Typical operating flow rates were 100 μL h^-1^ for the aqueous phase and 1000 μL h^-1^ for the oil phase in flow-focusing devices, and approximately 50 μL h^-1^ for each aqueous inlet and 1000 μL h^-1^ for the oil phase in co-flow devices. Flow rates were adjusted as required to obtain stable monodisperse droplets of the desired diameter. Emulsions were collected in microcentrifuge tubes for subsequent processing.

### Picoinjection

was performed using a dedicated microfluidic device to introduce the reaction mixture into pre-formed w/o droplets containing previously encapsulated and IPTG-induced *E. coli* DH10B cells. The picoinjection solution consisted of the screening assay reaction mixture lacking purified NanA, supplemented with lysozyme (4 mg mL^-1^, Roche), polymyxin B (8 mg mL^-1^, Sigma-Aldrich), and Texas Red-dextran (TexRed, 70 kDa, Invitrogen) at 0.1 or 0.25 mg mL^-1^ as a fluorescent tracer.

Picoinjection was achieved by electrocoalescence using integrated microelectrodes filled with 2.5 M NaCl and connected to a pulse generator (TGP110, AIM-TTi Instruments, UK) coupled to a high-voltage amplifier (Trek 2210, Trek Inc., USA). Electric pulses with a period of 50 μs, pulse width of 25 μs and amplitude of 150 V were applied to transiently destabilize the droplet interface and enable reagent injection. Flow rates were adjusted as required to maintain stable droplet spacing and uniform reagent delivery. Picoinjection was monitored using an inverted fluorescence microscope (DMi8, Leica Microsystems, Germany) equipped with a Phantom VEO 1310L high-speed camera (Vision Research, USA), which was also used during FADS experiments. Picoinjected droplets were collected for downstream processing.

### FADS

Simple monodisperse, co-flow generated, or picoinjected w/o droplets were analysed and sorted using a dedicated FADS microfluidic device. Depending on the experimental application, the platform was used either for fluorescence signal analysis or for the isolation of droplets containing putative NanA-positive clones.

For FADS experiments, the spacing and bias oil phases consisted of HFE-7500 fluorinated oil containing 1% and 0.5% (v/v) FluoroSurfactant, respectively. Flow rates were adjusted as required to maintain stable droplet spacing and reliable sorting. Droplets were interrogated using a fibre-coupled multi-wavelength laser module (CNI Laser, Changchung, China). TexRed fluorescence was excited at 594 nm and detected between 595-633 nm, whereas MH24 fluorescence was excited at 405 nm and detected between 420-470 nm.

Droplets exhibiting fluorescence within the predefined sorting gate triggered an electric pulse (500 V amplitude, 20 bursts at 25 kHz) applied through integrated microelectrodes, generating an electrostatic force that deflected positive droplets into the collection outlet while non-selected droplets were directed to waste. Sorted droplets were collected for downstream recovery and validation.

### Bacterial strains and cultivation

*E. coli* strains were routinely cultured in lysogeny broth (LB) medium at 37 °C with shaking at 180 rpm for approximately 12 h, unless otherwise stated. When required, LB medium was solidified with 1.5% (w/v) agar. Transformed clones were selected on LB agar supplemented with 100 μg mL^-1^ ampicillin.

## Supplementary Figures

**Supplementary Fig. S 1.**
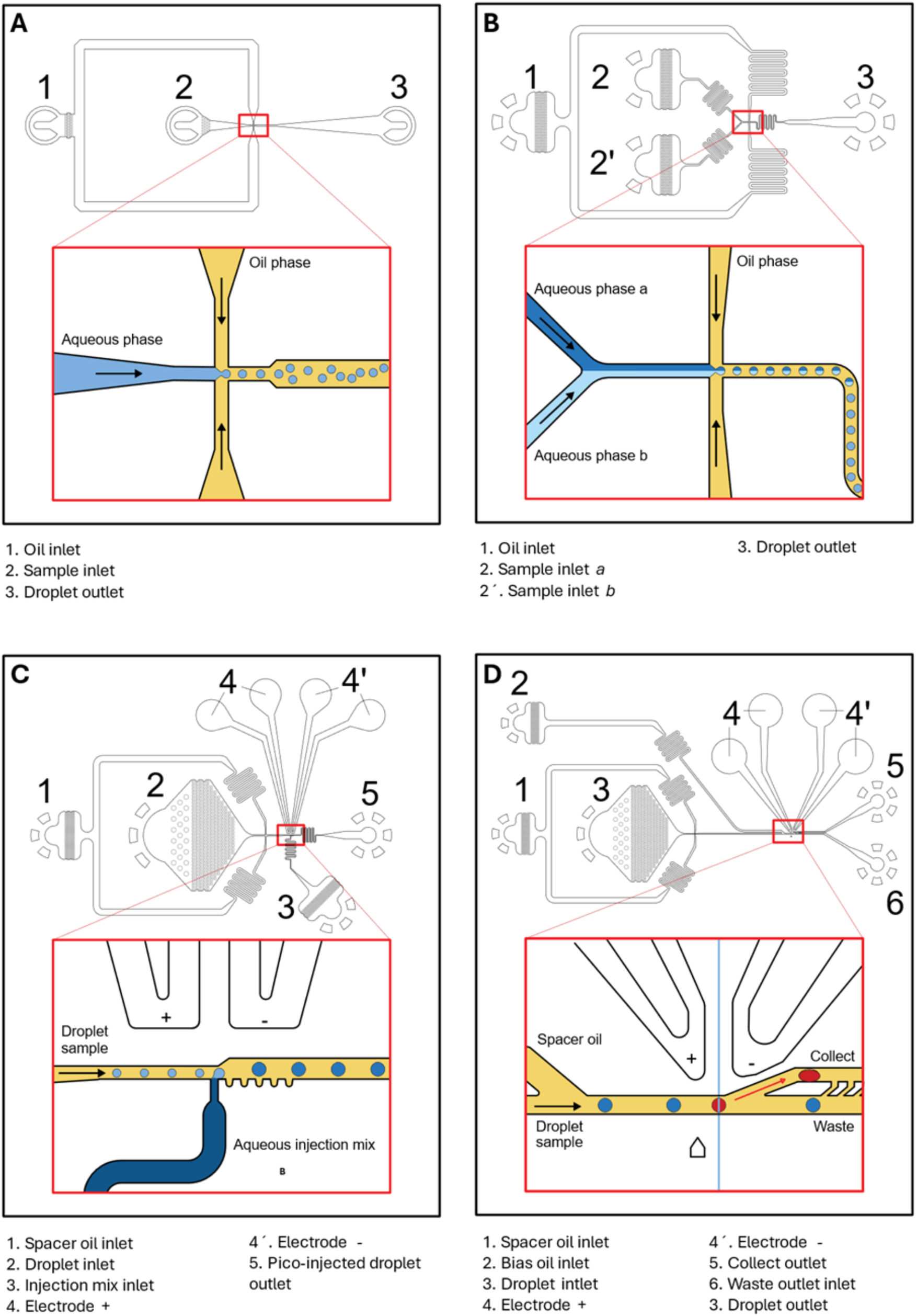
Designs of the four microfluidic device geometries used in this study. (A) Flow-focusing device for the generation of monodisperse w/o droplets from a single aqueous inlet. (B) Co-flow device enabling co-encapsulation of two aqueous phases in monodisperse w/o droplets. (C) Pico-injection device for the electro-triggered addition of aqueous reagents into pre-formed monodisperse w/o droplets. (D) Fluorescence-activated droplet sorting (FADS) device for electrostatic sorting of droplets based on fluorescence intensity.

**Supplementary Fig. S 2.**
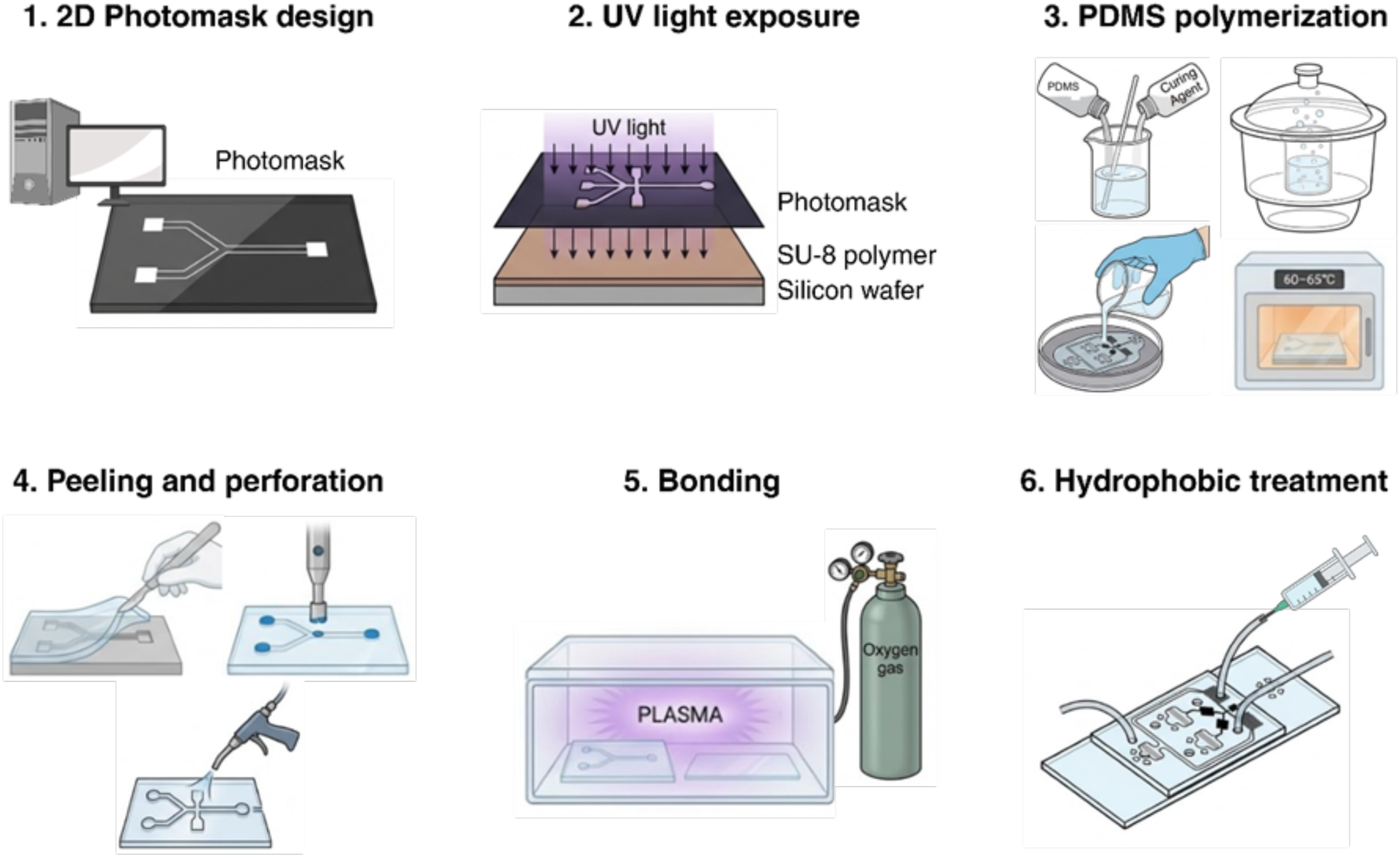
Step by step process for the fabrication of a microfluidic device. 1. 2D design of devices and photomask printing. 2. UV light exposition of photomask, SU-8 polymer and silicon wafer. 3. PDMS and curing agent mixing, degasification and polymerization. 4. Peeling, perforation and cleaning of the PDMS chip. 5. Exposition to oxygen plasma of PDMS and glass slide for covalent bonding. 6. Hydrophobic treatment of the device. Figure created with Biorender.

**Supplementary Fig. S 3.**
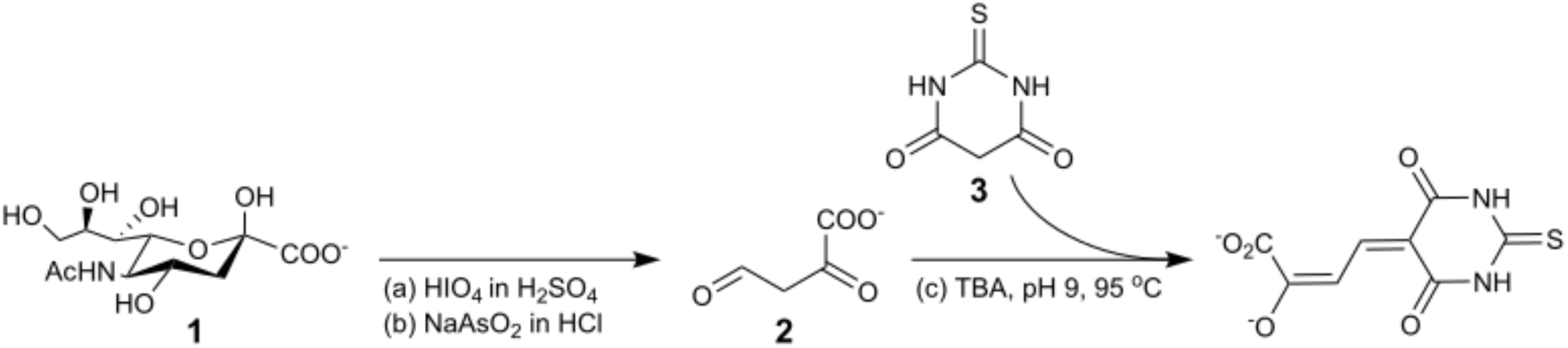
Schematic detection of TBA assay for the spectrophotometric detection of sialic acid. (1) N-acetylneuraminic aci), HIO4 (periodic acid), H2SO4 (sulphuric acid), NaAsO2 (sodium arsenate), HCl (hydrochloric acid), (2) 2-dioxobutanoate, (3) 2-thiobarbituric acid, TBA.

**Supplementary Fig. S 4.**
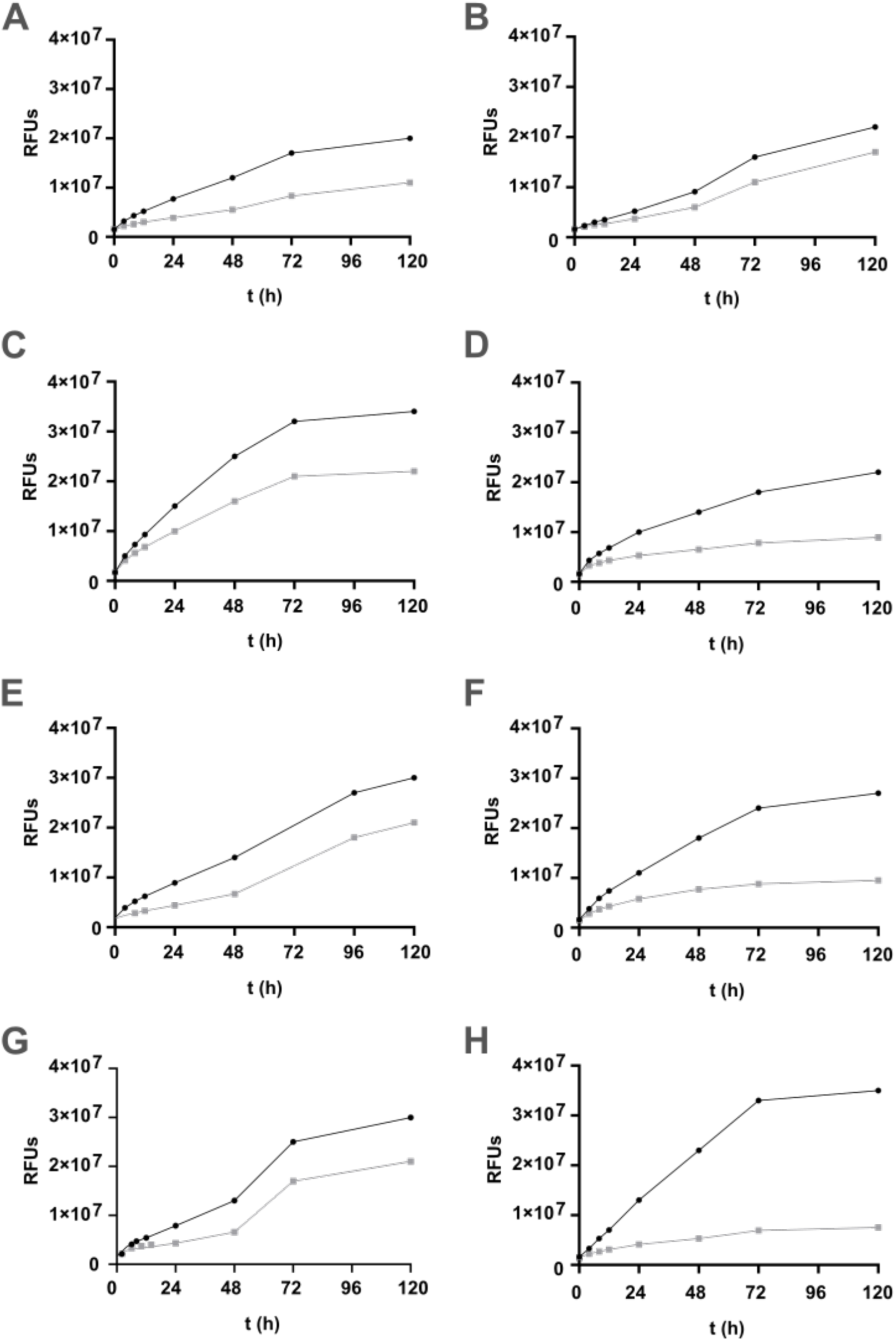
Monitoring of signal to background ratio between positive (black circles) and no-enzyme containing control reactions (grey squares). (A) CgKRED, (B) MoKRED, (C) MsKRED, (D) AmKRED, (E) AsKRED, (F) SsKRED, (G) AbKRED, (H) FbKRED. Data show endpoints (n=3).

**Supplementary Fig. S 5.**
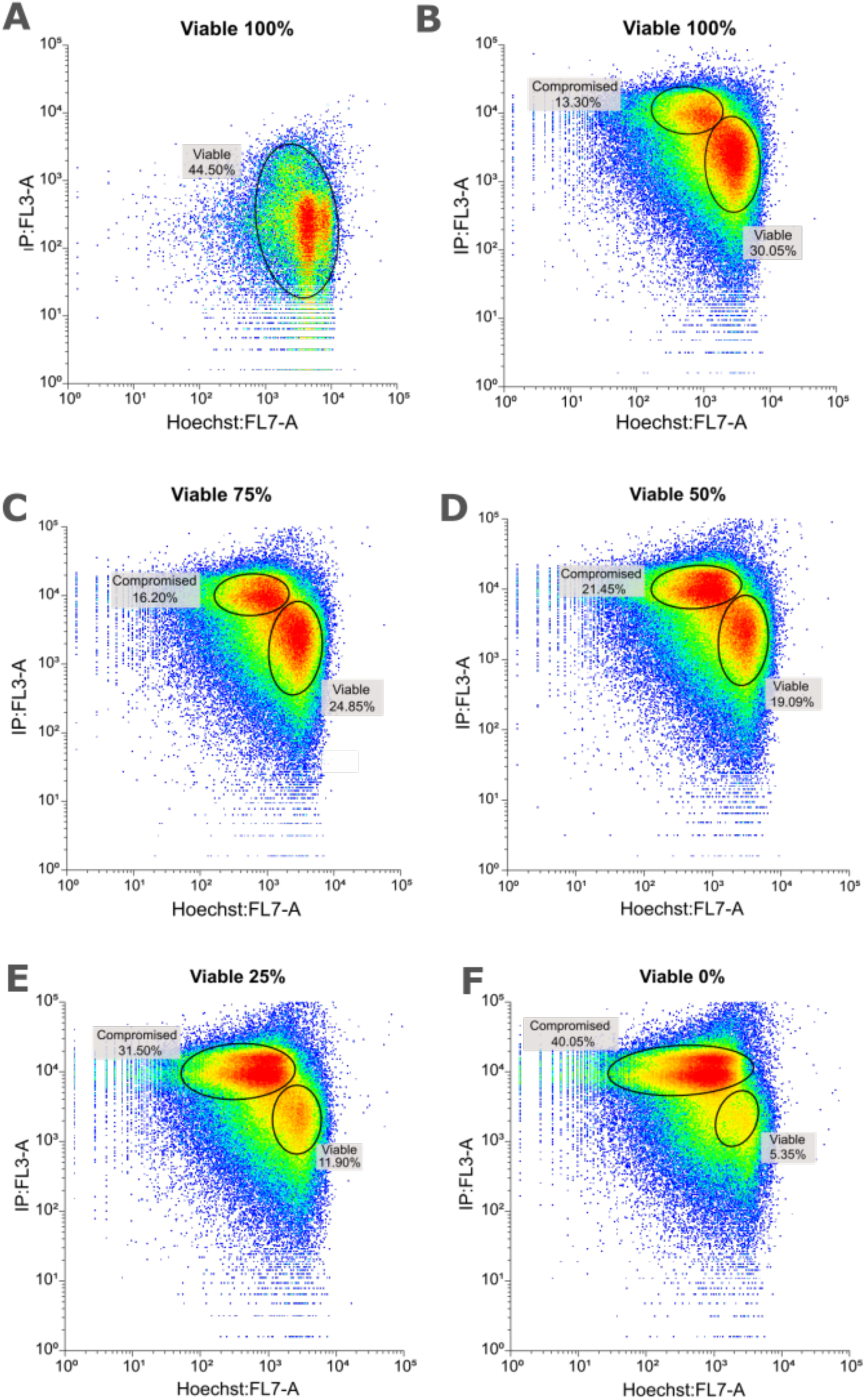
Flow cytometry analysis of the calibration method using Hoechst 33342 (H33342, x-axis) and propidium iodide (PI, y-axis). (A) Untreated cells stained with H33342 only. (B) Untreated cells stained with both dyes. (C-F) Mixtures of untreated and heat-thaw treated cells (75/25, 50/50, 25/75 and 0/100 viable/compromised). Images created with Floreada.io

**Supplementary Fig. S 6.**
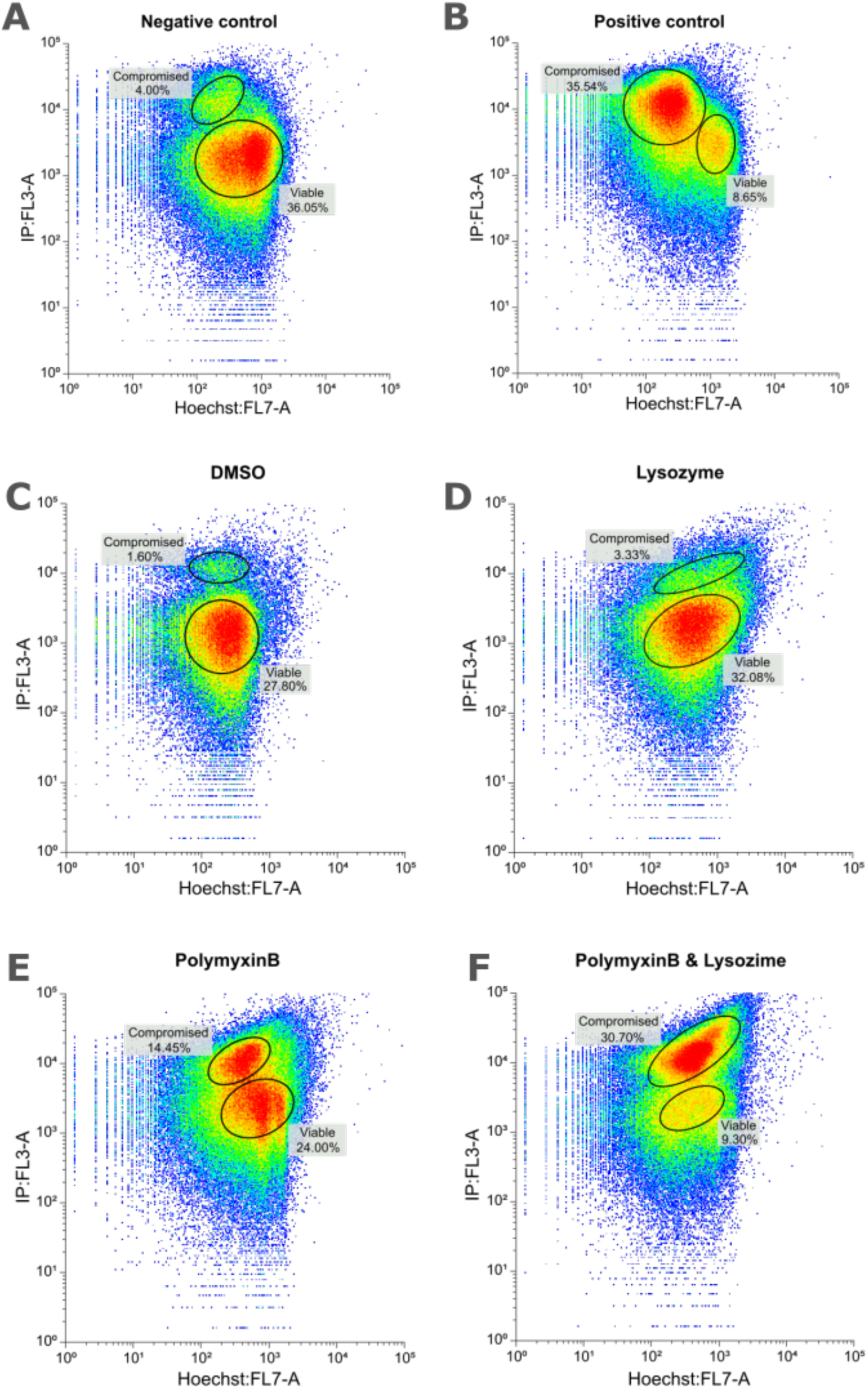
Flow cytometry analysis of membrane permeability treatments using Hoechst 33342 (H33342, x-axis) and propidium iodide (PI, y-axis). Untreated negative control (a), heat-thaw treated positive control (b), 2% DMSO (c), lysozyme (d), polymyxin B (e) and lysozyme plus polymyxin B (f) are shown. Images created with Floreada.io

**Supplementary Fig. S 7.**
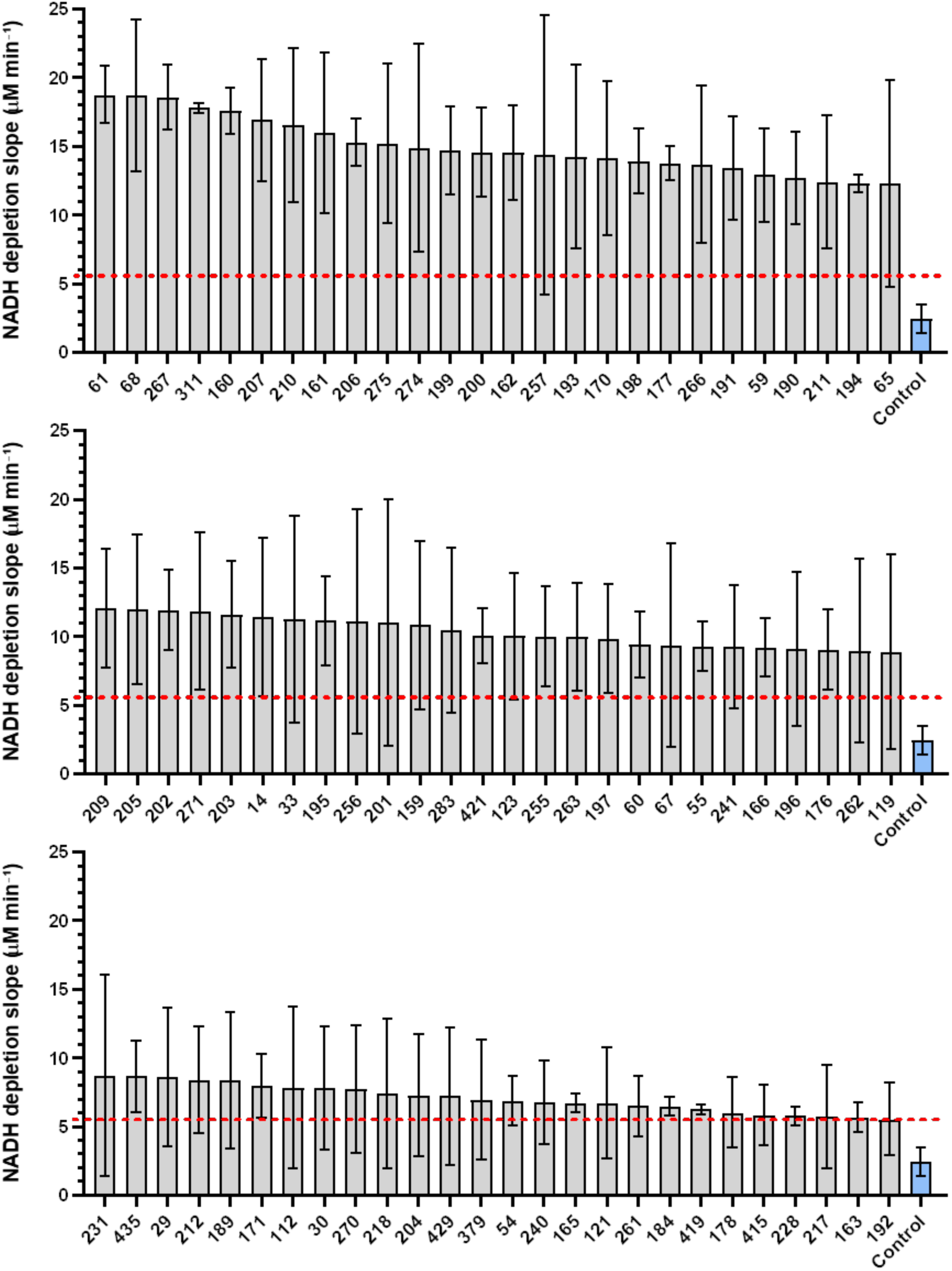
Bar graphs showing the mean NADH depletion slopes as measurement of the enzymatic activity for the 78 selected metagenomic clones. Red lines indicate the activity cut-off threshold at 5.57 μM min⁻¹. Numbers on the X axis indicate number given to each clone. Control in light blue corresponds to the negative control (2.44 μM min⁻¹) consisting of the empty pBS CFE.

**Supplementary Fig. S 8.**
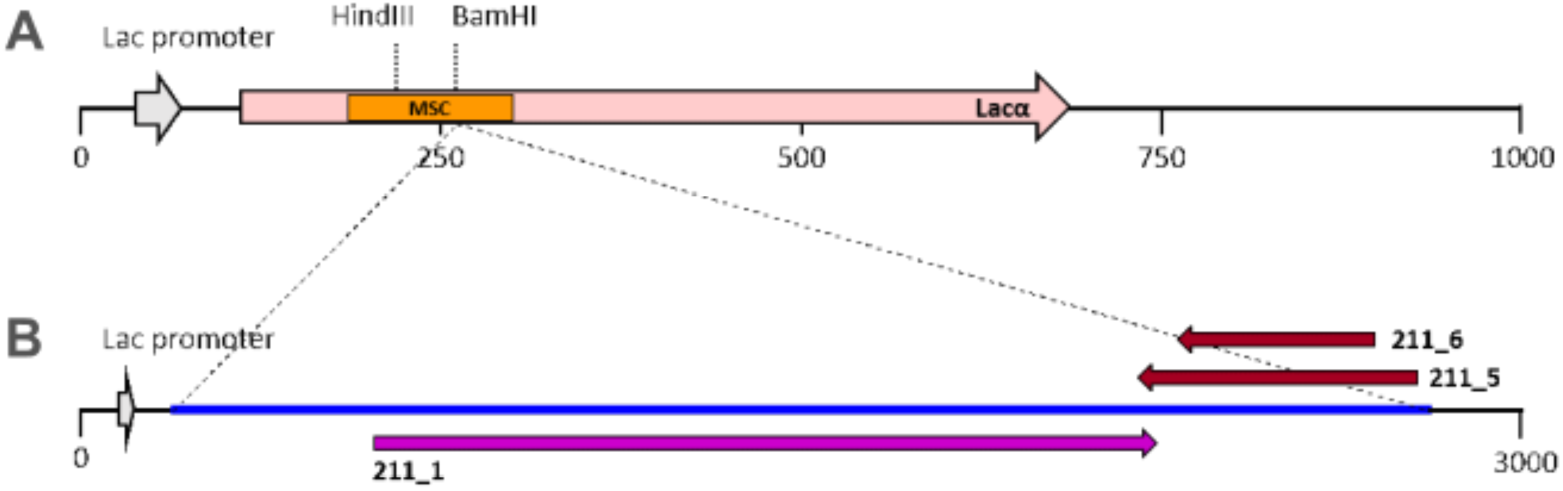
Schematic representation of metagenomic insert 211 cloned into pBluescript SK (+). Black lines represent the vector backbone, while the blue line represents the metagenomic insert. The Lac promoter is shown as a grey arrow. Predicted ORFs are indicated by arrows: pink denotes lacZα, purple indicates the ORF oriented in the same direction as the Lac promoter, and red indicates ORFs oriented in the opposite direction. The multiple cloning site (MSC) is shown in orange. BamHI indicates the cloning site, while HindIII is shown for mapping purposes only. Numbers on black and blue lines represent nucleotide coordinates (bp), and labels next to arrows represent ORF identifiers. Panels show (A) the pBluescript SK (+) vector and (B) clone 211.

## Supplementary Tables

**Supplementary Table S 1.** Plasmids used throughout this study.

| Plasmid | Description | Application |
| --- | --- | --- |
| pkk223-3 | AmpR, tac promoter | Expression of recombinant NanA by native RNA polymerase |
| pBluescript SK (+) | AmpR, lacI promoter, T7 promoter, T3 promoter, lacZ-α fragment | Cloning and amplification, high copy number vector. Enables white/blue screening of clones with lacZα system |

**Supplementary Table S 2.**
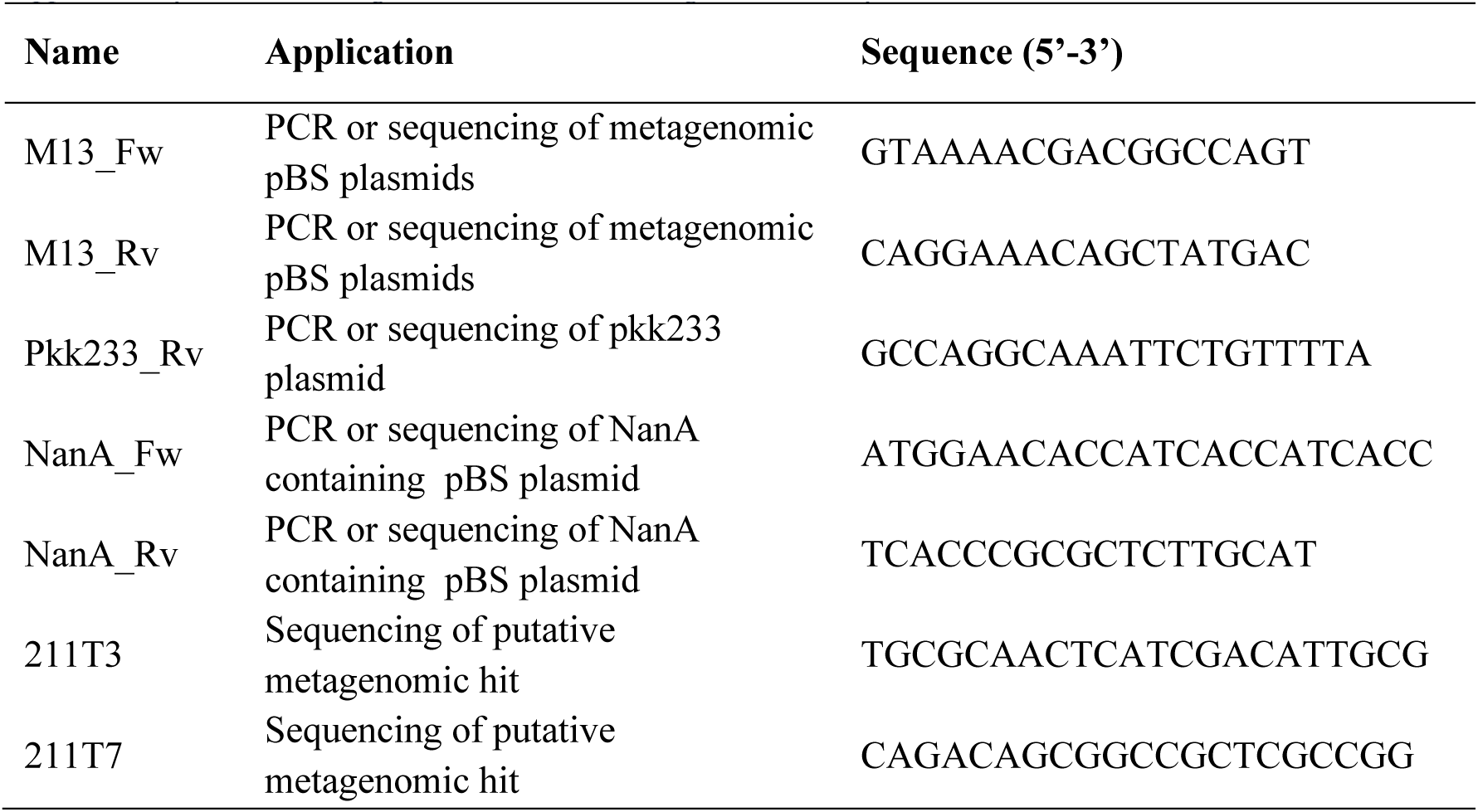
Oligonucleotides used throughout this study.

**Supplementary Table S 3.** E. coli strains used throughout this study.

| Strain | Genotype | Application |
| --- | --- | --- |
| DH5α | F- end A1 glnV44 thi-1 recA1 relA1 gyrA96 deoR nupG purB20 ϕ80dlacZ M15 $\Delta$ (lacZYAargF) U169, hsdR17 (rK-mK<sup>+</sup>), $\lambda^-$ | Molecular cloning and amplification |
| BL21 (DE3) | F- ompT gal dcm lon hsdSB (rB<sup>-</sup> mB<sup>-</sup>) $\lambda$ (DE3 [lacI lacUV5-T7p07 ind1 sam7 nin5]) [malB<sup>+</sup>] K-12 (S) | Protein expression |
| ElectroMaxTM DH10B | F- mcrA $\Delta$(mrr-hsdRMS-mcrBC) ϕ80lacZΔM15 ΔlacX74 recA1 endA1 araD139 $\Delta$ (ara, leu)7697 galU galK λ-rpsL(StrR) nupG | Library construction, transformation efficiency, DNA recovery |

**Supplementary Table S 4.**
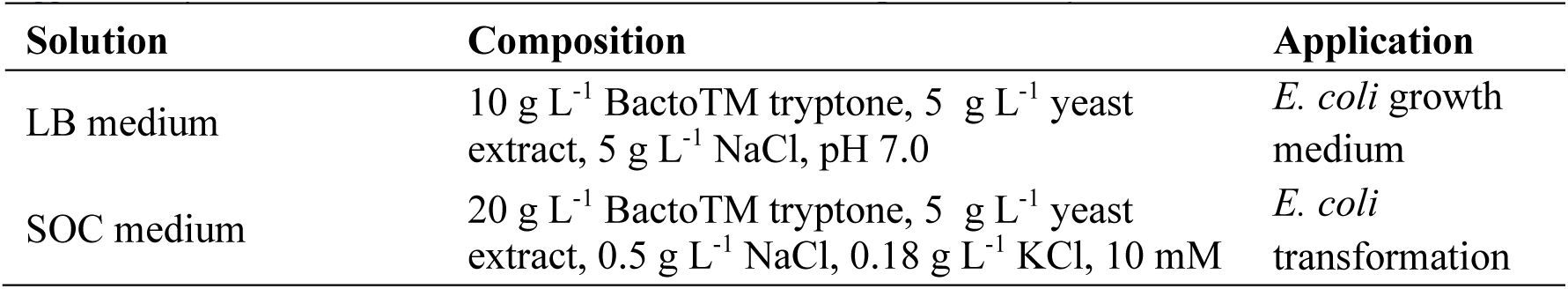
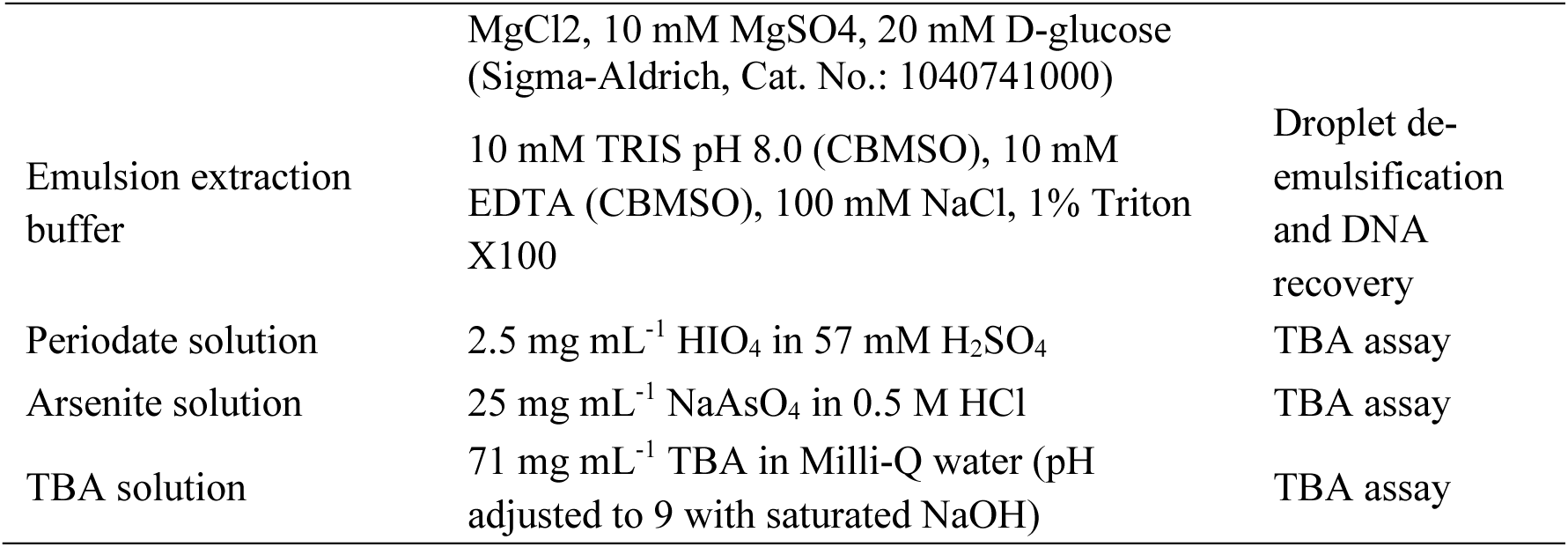
Culture media and solutions used throughout this study.

**Supplementary Table S 5.**
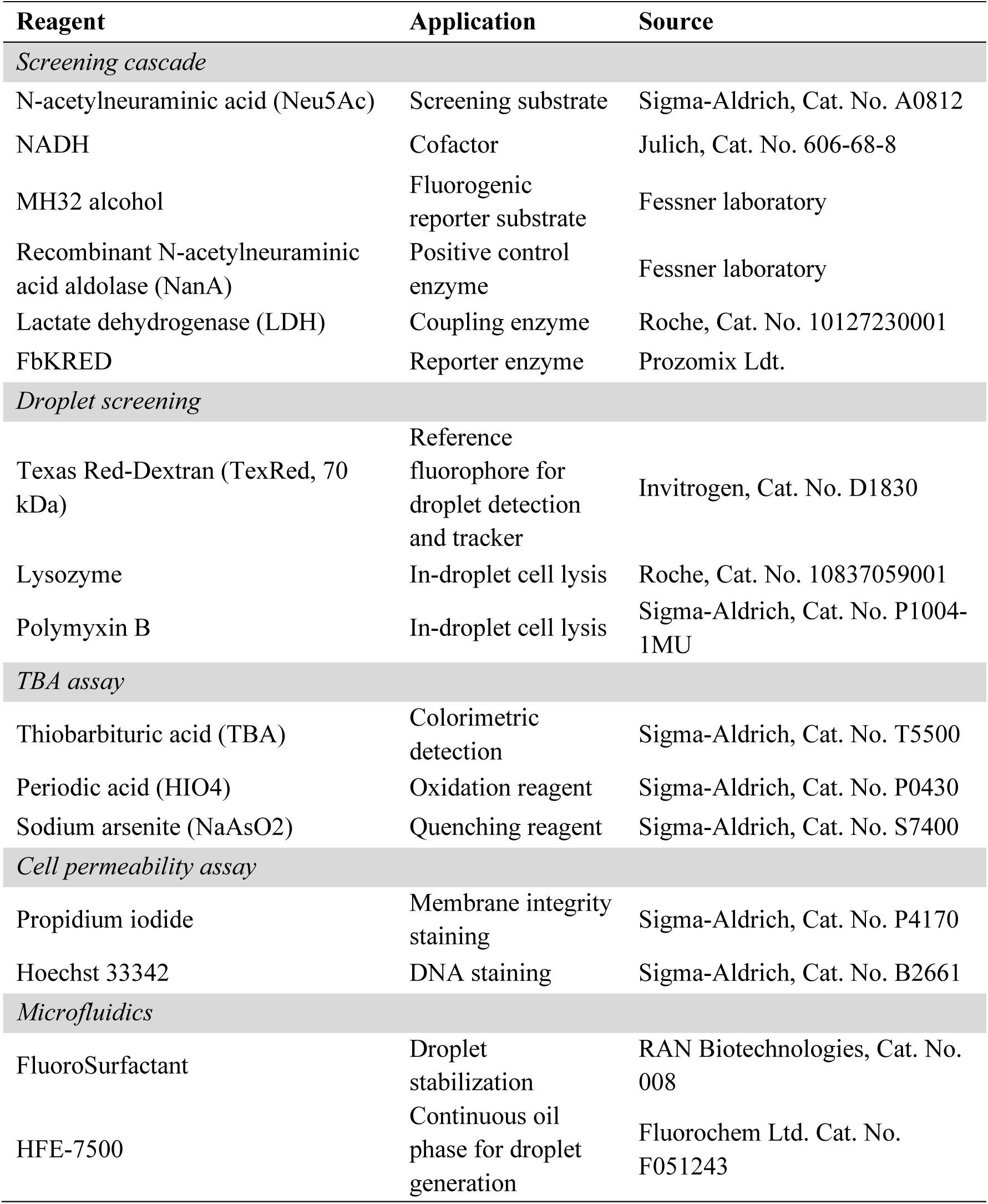
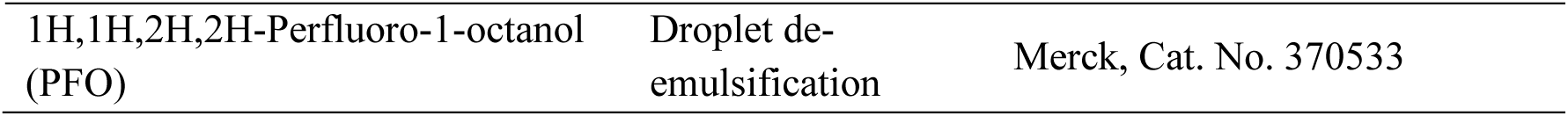
Key reagents used throughout this study.

**Supplementary Table S 6.** Characteristics of the predicted ORFs.

| ORF | BLASTP | InterPro |  |  | Pyre2 |
| --- | --- | --- | --- | --- | --- |
|  |  | Family | Domain | Superfamily |  |
| 211_1 | β-(1-6) glucan synthase. Coverage: 100% Identity: 92% | Glucan 1,3-β-glucosidase | Glycoside hydrolase 17 | Glycoside hydrolase, Transglycosylase | β-1,3-glucanosyltransferase |
| 211_5 | Threonine aldolase. Coverage: 99% Identity: 97% | - | β-eliminating lyase | PLP-dependent transferase | PLP-dependent transferase |
| 211_6 | Threonine aldolase. Coverage: 64% Identity: 66% | - | - | - | - |

## Sequence ORF 211_1

MVAAVWWWLATPVALTRAPIDGAAKLECVSYAPFRGDQTPHDPTLIVAPAQIAEDLAELANVSKCIRTYSIDNGLDKVPELASRVGLKVLLGVWIGRDRARNAQLIDIAVSLVKDHPGTIAALIVGSEVLLRGDMTVGDLRDTIRSVRPRVDVPVTYADVWEFWLRYREVGADVDFVTAHFLPYWEDVPPRAEDAAAHVDGYRKQVVAAFPGKEILIGETGWPSHGRMRDGALASRVNQARFISEILERARRDNFRVNLFEAYDEPWKRRWEGTVGGYWGLFDGAERKLKYPAGVAISNYPLWKLQMACGMLLCIGIFATALFTLKRRPSPTPLASWLAVAVSATLGGILLGVSADKMLHESYGLGGWLVQGFLLAAAVMAPLLSTYALMSGRALPAFLEVLGPSKGLTPLFMSNMLGITLIVTTLIAAQTALSLIFDARWRDFPFAALTMAVVPFWTLAFLNGSKSGERPLSEAVFAGLFALAAIYVVFNEGFENWQAMWTGAIYVVLGSALWRARTAAVA*

## Sequence ORF 211_5

VLDEVVAIAKANGLITHMDGARLLNACVATKISAKDMAAGWDSTWIDFSKGLGAPIGGVIAGSRAFIDDVWRWKQRLGGSMRQAGIAAAACVYALDHHVDRLADDHANARALARGLSQINGIEVQEPETNLVFFKPDGAGIAGDKMVEALRKRGVLLAMMDGRIRACTHLDVRADMIEETIGIVREIVRGA*

## Sequence ORF 211_6

MGFDLDRFLKRPRRADRRRDCGIARLHRRRLALETAARRIDAAGRHCCGRLRLRARPSRRPPRRRSRQCAGAGARAVADQRHRSARARDQSGVLQARWRRYRRRQDGRGLAQARRPARDDGRPHSRLHPSRRPR*

